# Naturally occurring Alzheimer’s disease in rhesus monkeys

**DOI:** 10.1101/2022.10.20.513120

**Authors:** Zhenhui Li, Xiaping He, Shihao Wu, Rongyao Huang, Hao Li, Zhengbo Wang, Limin Wang, Dongdong Qin, Yu Kong, Yingqi Guo, Xia Ma, Christoph W. Turck, Zhiqi Xiong, Wenchao Wang, Xintian Hu

## Abstract

Alzheimer’s disease (AD) is the most common neurodegenerative disease. To date, its cause is unclear and there are no effective treatments or preventive measures. Despite there are accumulating evidences for the existence of AD pathological hallmarks in the brain of aging rhesus monkeys, it remains a mainstream notion that monkeys do not develop AD naturally. This is an important issue because it will determine how we use monkeys in AD studies. To settle down this issue, a group (n=10) of aged rhesus monkeys 26 years old or above went through a systematic AD screening procedure in this study. Three of these monkeys showed severe memory impairments (SMI) after evaluated with a classic working memory test. Further behavioral testing revealed that the SMI monkeys also exhibited apathy-like behavior, which is another core AD clinical symptom. In addition to the cognitive deficits, two of the three SMI monkeys developed all of the three AD pathological hallmarks, including neurofibrillary tangles, senile plaques and neuronal loss. According to the diagnostic criteria of human AD, the two SMI monkeys were clearly naturally occurring AD monkeys. These results suggest that AD is not a uniquely human disease and monkeys have great potential for the development of much needed etiological AD models, which are vital for better understanding of developmental process of AD and the base of identification of early diagnostic biomarkers and effective therapeutic targets of AD.

## 1. Introduction

Alzheimer’s disease (AD) is the most common neurodegenerative disease (*1, 2*). It usually afflicts people over 65 years old and the morbidity increases dramatically with age (*3*). AD is always accompanied with symptoms of conspicuous cognitive degeneration, including short-term and long-term memory impairments, apathy, attention-deficits and visuospatial function impairments (*4-6*). In addition to accumulation of protein fragment β-amyloid (Aβ) outside neurons to form senile plaques (SPs), and hyperphosphorylated tau protein clumping into long filaments that twist around each other inside neurons to form neurofibrillary tangles (NFTs), typical AD pathological hallmarks also include significant neuronal loss.

Over past decades, many important AD discoveries have been made from studies based on transgenic mouse AD models, but the therapeutic treatments developed on these models are almost untranslatable to AD patients (*7-11*). This failure has been partially attributed to the fact that rodents and human are far apart in the evolution, and as a result, the structure, function and metabolism of their central nervous system (CNS) are quite different from each other. On the other hand, rhesus monkeys, which are evolutionary close to human, have relatively longer-life span, highly developed CNS, higher levels of cognitive functions, more complex behaviors than those of rodents and, therefore, are a much-preferred species for developing AD models since AD is a cognitive deficits based brain disease. In particular, non-human primates have been shown to spontaneously develop brain diseases previously thought to be unique to human beings, including recent two reports from our lab on the first case of monkey with naturally occurring Parkinson’s disease (*12, 13*) and first two cases of monkeys with spontaneous autism spectrum disorders (Preprint, https://doi.org/10.1101/2022.03.17.484827). However, whether monkeys can develop AD spontaneously is undetermined. One important but usually ignored fact is that rodents do not suffer from AD naturally (*14*). If monkeys do not develop AD naturally either, there is probably no fundamental difference of using monkeys to develop AD model compared with rodents. Therefore, this need to be clarified before we can conclude with high confidence that monkeys are ideal experimental animals in translational AD studies.

As the first step of finding spontaneous AD monkeys, there are several studies on whether monkeys can spontaneously develop AD pathological changes in their brains. Since Wisniewski’s first study in 1973 (*15*), it has been generally accepted that monkeys can spontaneously develop SPs in their brains as they aging (*16-18*). But the evidence for monkey tau pathology had been missing till recently. In 2017, using immunoelectron microscopy technology, Paspalas and colleagues provided strong evidence for the existence of NFTs in an aged rhesus monkey brain accompanied by SPs (*19, 20*). However, it is unknown whether the appearance of SPs and NFTs in the monkey brain was associated with cognitive impairments, which are the core clinical symptoms of AD. Autopsy studies revealed that even though some aged people had severe SPs and NFTs pathology in their brain, their cognition had been normal before death and, thus, they were not AD patients (*21*). Therefore, if monkeys who have SPs and NFTs pathology are not accompanied by cognitive impairments, we still cannot conclude that they are AD monkeys. Previous studies focused on whether monkeys could naturally develop AD pathology, and, in this approach, the monkeys had been sacrificed to perform the pathological tests before any cognitive tests could be carried out. This limitation makes this strategy unable to directly and effectively answer the question: are there naturally occurring AD monkeys?

How can we find the direct answer then? In 1906, Dr. Alzheimer reported the first AD case: a 51-year-old female patient whose main clinical symptoms were cognitive disorders such as impaired working memory, aphasia and poor directionality. After the autopsy, he found that there were many amyloid plaques (which have been named later as senile plaques) in her brain tissues, entangled structures (which have been named later as neurofibrillary tangles) in the neurons, and severe atrophy of the cerebral cortex. Although the etiology and development mechanism of AD are still unclear, the standard of defining AD by combining typical clinical symptoms with classical pathological hallmarks used by Dr. Alzheimer has been generally accepted, and confirmed recently (*22, 23*). Therefore, in order to answer the question of whether rhesus monkeys can spontaneously develop AD, it is necessary to look for aged monkeys with cognitive impairments first, and then to investigate whether these impairments are accompanied by classical AD pathological changes, including SPs, NFTs and neuronal loss. Therefore, in this study, our work started from behavioral phenotype screening and then go to pathological investigation, just the opposite of previous studies.

## 2. Methods

### 2.1. Animals

The experimental protocol had been approved by the Ethics Committee of Kunming Primate Research Center (Approval ID: IACUC18001) affiliated to Kunming Institute of Zoology, Chinese Academy of Sciences.

For the huge workload of cognitive assessment of monkeys, we picked the 10 oldest rhuses monkeys (*Macaca mulatta*, 7 males, 3 females, over 26 years old) from a group of 42 aged population, all over 19 years old from the Kunming Primate Research Center to perform cognitive tests. The monkeys (Table 1) were moved into individual cages in monkey rooms with a standard 12 h light/ 12 h dark cycle (light on at 07:00 a.m.). They were fed with commercial monkey biscuits twice a day, and fruits or vegetables once a day with tap water *ad libitum*. For their old ages, the monkeys were under careful veterinary oversight to ensure a good health. Best efforts were made to minimize the number of monkeys and their suffering involved in this study. Monkeys were treated in accordance with the National Institute of Health (U.S.A) Guide for the Care and Use of Laboratory Animals (8^th^, edition, 2011; National Academies Press, Washington, DC) as well.

**Table 1.** Information of the ten aged monkeys in this study and their Maximum B Values.

| Groups | Subject | Age<sup>§</sup> | Gender | Maximum B Value (s) | Grouped B Value Mean±SEM (s) |
| --- | --- | --- | --- | --- | --- |
| Normal aged monkeys | 1<sup>&#</sup> | 29 | M | 38 | 36±0.63 |
|  | 2<sup>&#</sup> | 27 | M | 37 |  |
|  | 3<sup>&</sup> | 26 | F | 36 |  |
|  | 4<sup>#</sup> | 29 | M | 35 |  |
|  | 5<sup>&</sup> | 26 | M | 35 |  |
| Mild Memory Impairment (MMI) monkeys | 6 | 28 | M | 16 | 15±1 |
|  | 7 | 27 | M | 14 |  |
| Severe Memory Impairment (SMI) monkeys | 8<sup>#</sup> | 28 | M | 8 | 7±0.58 |
|  | 9<sup>&#</sup> | 27 | F | 7 |  |
|  | 10<sup>&#†</sup> | 26 | F | 6 |  |

§: age at the time of sacrifice; &: the monkey also tested with HIT; #: monkeys that sacrificed for pathological test; †: the monkey that underwent a biopsy for immunoelectron microscopy. Note that to minimize the number of sacrifice animals, only the three SMI monkeys (most likely to develop AD pathology) and three normal aged monkeys (as pathological controls) were sacrificed for the pathological tests.

### 2.2. Experiment overview

In this study, the first step was to access AD candidate monkeys’ cognitive functions. Because age is the greatest risk factor for AD development (*24*), ten oldest rhesus monkeys aged 26 or above were selected to perform the Variable Spatial Delayed Response Task (VSDRT) to evaluate their working memory function, and to perform the Human Intruder Test (HIT) to assess their emotion function. The results demonstrated that the 10 monkeys could be divide into three groups: normal aged (n=5), mild memory impairment (MMI, n=2) and severe memory impairment (SMI, n=3) monkeys (see results). Then the three SMI monkeys who were most likely to possess AD pathology, and three normal aged monkeys (as controls for AD pathology) were sacrificed to carry out pathological validation. In the tests, immunohistochemistry and immunofluorescent staining were performed to explore the presence of NFTs and SPs in the SMI monkeys’ brains. In addition, immunoelectron microscopy was employed to illustrate the submicroscopic structures of the NFTs identified by the staining. Finally, apoptotic cells were illustrated by terminal deoxynucleotidyl transferase (TdT) labeling to demonstrate the cell death in the brain regions that are associated with the pathogenesis of AD.

Details of the above tests are described in following sections.

### 2.3. Cognitive tests

#### 2.3.1. Variable Spatial Delayed Response Task

The Variable Spatial Delayed Response Task (VSDRT) has been widely used to measure the capacity of visuospatial working memory of the brain (*25-28*), a function that is severely impaired in AD patients (*29*). During the test, the monkey was placed in a Wisconsin General Test Apparatus (WGTA) situated in a sound-attenuated room, and was always tested at the same time of the day immediately prior to feeding (*30*).

The VSDRT consisted of two stages, a training stage and a testing stage. During the training stage, the monkey was initially trained for a 2-well spatial delayed response task for 1000 trials. At the beginning of each trial, the monkey watched the experimenter placing a piece of food bait in 1 of the 2 food wells on a wooden board located in front of the monkey as a reward. The two food wells were then covered with two identical wooden planks, and an opaque screen was pulled down between the monkey and the food wells for a specified delay time period. At the end of this delay, the screen was raised up and the animal was allowed to choose freely the reword from one of the two wells. The reward was quasi-randomly distributed between the left and right wells over 30 trials that made up a daily training session. The delay time period was increased or decreased depending on whether the animal had a correct response rate greater than 90% (27 correct out of the 30 trials). At the end of the 1000-trial training stage, the monkey had fully mastered the task.

During the testing stage, in order to identify the capacity of each monkey’s memory, the animal was further trained for a variable delayed response task, in which the delay period was varied quasi-randomly for five different delay time periods (namely A, B, C, D and E; A=0×B; B=1×B; C=2×B; D=3×B; E=4×B) for a total of 30 trials in a daily test session. The beginning B value depended on the training stage: if the monkey’s performance was not high during the training stage, the B value in the testing stage started at 1 second. Otherwise, the B value started at a higher number, which was decided by the experienced trainer. Until the animals achieved a stable performance of 90%-99% correct for 3 consecutive days, the B value was increased by one second per step. If the accuracy reaches 100%, the B value was increased by one second in the next session. If the performance was less than 90% on a certain B value for 3 consecutive days, the B value was reduced by one second. We used the maximum B value that the monkey could achieve stably for at least 5 consecutive days of 90% correct or above to represent its memory capacity.

#### 2.3.2. Human Intruder Test

The Human Intruder Test (HIT) is a widely used non-invasive test to assess a monkey’s responses to changes in the environment in a laboratory setting (*31, 32*). This test poses only a mildly threatening stimulus and a range of emotional reactions can be observed.

The HIT used in this study consisted of a 10-minute baseline phase (camera-only), and of which the last two minutes were used for behavioral analysis, followed by three 2-minute intruder phases. During the intruder phases, a person unfamiliar to the monkey stood 60 cm in front of the monkey’s cage in three orientations in succession. The stranger initially stood orthogonal to the subject (profile phase), then turned to directly face the subject (stare phase), and then turned directly back to the subject (back phase). After the back phase, the intruder left the room and the test was completed. The intruder used a timer to start and end each phase.

The intruder for all the monkeys was the same person. All the video clips of the tests were scored using a standard second-by-second analysis protocol (*32*). Briefly, each video clip was scored independently by three experienced observers, who were blind to the conditions of the test. The last two minutes of the baseline phase and the three two-minute intruder phases were scored for the duration of nine behaviors: staying at the back of cage (at least three limbs occupy the back half of the cage), pace (repeated locomotion for 3 or more times), freeze (no movement), lipsmack (the mouth was puckered and moving quickly up and down to produce a smacking sound, often paired with eyebrows and ears drawing back), teeth gnash (a chewing motion of the mouth with no food or objects involved), fear grimace (grin-like facial expression with lips drawn back showing clenched teeth, often paired with flapping of ears and stiff body posture), yawn (a slow opening of the mouth to an extremely wide position, often exposing the teeth), scratch (a vigorous stroking of the body with nails), and threating/cage shaking (staring at the intruder with open mouth, eyebrows lifted, ears flattened or flapping, rigid body posture, lunging toward the front of the cage, shaking cage vigorously, slapping cage). After the data analysis, only the staying at the back of cage and threating/cage shaking were included in the further detailed analyzing data set, for these two behaviors were the dominant reactions of all the tested monkeys and the only two displayed by all of them. The staying at the back of its cage represented the monkey’s mild reaction (a low reaction) to the intruder (*33*), and the threating/cage shaking was an exaggerated aggressive response to the intruder, representing a high reaction (*34*). These two behaviors to some extent antagonize each other since they were the dominant ones: more of one means less of the other. If staying at the back of its cage increases, the threating/cage shaking will decrease, which indicates the animal show an apathy tendency. The differences of the two behaviors between the normal aged monkeys (Monkey 1, Monkey 2, Monkey 3 and Monkey 5) and the SMI Monkey 9 and 10 were compared. When the HIT experiment began, the normal aged Monkey 4 (29 years old) and SMI Monkey 8 (28 years old), two of the oldest individuals in the group, became very weak. Considering the importance of the next pathological experiments and to ensure the quality of the HIT data, they were sacrificed without taking the HIT test.

### 2.4. Pathological tests

#### 2.4.1. Brain acquirement, fixation and section

After the behavioral tests had been completed, three normal aged monkeys (Monkey 1, 2 and 4) and three SMI monkeys (Monkey 8, 9 and 10) were anesthetized with ketamine (10 mg/kg, *i.m*.), followed by euthanasia with sodium pentobarbital (100 mg/kg, *i.m*.) and trans-cardiac perfusion with PBS. Each brain was removed from the skull and cut into left and right hemisphere. The right hemisphere was immersed into 4% paraformaldehyde (PFA), 0.01 M PBS at 4 °C for 1 week. After that, the frontal lobe, parietal lobe, occipital lobe and temporal lobe, hippocampus and entorhinal cortex of each right hemisphere were dissected (Figure. S1), cutting into 3 mm-thick sections and embedded in paraffin or 30% agar. The paraffin embedded sections were then sliced into 4 μm-thick slices on a Leica RM2016 Microtome (Leica, Shanghai, China) for immunohistochemistry staining and apoptotic cells labeling. The agar embedded sections were sliced into 40 μm-thick free-floating sections slices on a Leica VT1000 S Vibratome (Leica, Shanghai, China) for immunofluorescent staining. The left hemisphere was blocked to 50 different regions (covering the cerebral cortex, basal ganglia, limbic system, diencephalon, cerebellum and brainstem) and stored at −80 °C for future molecular studies. All the brains did not show notable ischemic lesions or tumors.

#### 2.4.2. Antibodies

The primary antibodies used for the immunohistochemistry or immunofluorescence staining and immunogold labeling were 6E10 (EMD Millipore, Cat. #MAB5206), GFAP (Abcam, Cat. #ab7260), Iba1 (Wako, Cat. #019-19741), AT8 (Thermo Scientific, Cat. #MN1020), AT100 (Thermo Scientific, Cat. #MN1060) and NFTs (EMD Millipore, Cat. #AB1518). The secondary antibodies for the immunohistochemistry or immunofluorescence staining and immunogold labeling were goat anti-mouse IgG (ABcam, Cat. #ab6789), Cy2 AffiniPure donkey anti-rabbit IgG (Jackon ImmunoResearch, Cat. #711-225-152), Cy3 AffiniPure donkey anti-mouse IgG (Jackon ImmunoResearch, Cat. #715-165-150), goat-anti-mouse conjugate to 10 nm gold (Sigma, Cat. #G7777) and goat-anti-rabbit conjugate to 20 nm gold (abcam, Cat. #ab27237).

#### 2.4.3. Immunohistochemistry staining

For the immunohistochemistry staining, the paraffin sections were mounted on microscope slides and baked at 65 °C overnight. Then, the sections were deparaffinized and rehydrated with two 10-minute changes of xylene and graded alcohols, respectively. The treated sections were later heated for 2 min in citrate buffer in a pressure cooker for antigen retrieval. Then the sections were treated with 70% formic acid for 7 min. Endogenous peroxidase of the sections were inactivated by incubation with 3% hydrogen peroxide in methanol for 10 min. The sections were then blocked with 2% bovine serum albumin (BSA) in 0.5% Triton X-100 for 30 min at 37 °C. The primary antibodies were diluted in blocking solution and incubated with the sections at 4 °C overnight. The sections were then treated with HRP-conjugated goat anti-rabbit/mouse secondary antibodies at room temperature for 2 h and developed using 3, 3’-diaminobenzidine (DAB, MXB, China) in chromogen solution, and counterstained with hematoxylin (Sigma, USA) for cell nucleus identification. In the negative-control experiments, only the primary antibodies were omitted. Last, the sections were dehydrated and cleared with graded alcohols and xylene, respectively, and then covered with neutral gum and cover slips. The slides were analyzed using an Olympus CX41RF microscope (Olympus, Tokyo, Japan) and the images were captured with the matching Olympus DP25 microscope digital camera.

#### 2.4.4. Immunofluorescent staining

The free-floating sections were used for immunofluorescence staining. Before the staining, autofluorescence was quenched by incubation with 0.06% potassium permanganate for 10 min at room temperature. All the free-floating sections were then blocked in 2% BSA in 0.5% Triton X-100 for 1 h at room temperature. The primary antibodies were diluted in blocking solution and the sections were incubated at 4 °C overnight, followed by incubation with the corresponding fluorescent secondary antibodies at room temperature for 2 h. Nuclei were stained with 4’,6-diamidino-2-phenylindole (DAPI, Sigma, USA) for 15 min. In the negative-control experiments, the primary antibodies were omitted. The slides were mounted with Fluoromount-G (SouthernBiotech, USA) and imaged on a laser scanning confocal microscope (A1, Nikon, Japan).

#### 2.4.5. Apoptotic cells labeling

The deparaffinization and hydration of the paraffin sections in the Apoptotic cells labeling process were similar to those in the immunohistochemistry staining. The paraffin sections were assayed for apoptotic cells using the ApopTag Plus Fluorescein *in Situ* Apoptosis Detection Kit (Millipore, Cat. #S7111) as per the manufacturer’s instructions. The slides were mounted with Fluoromount-G (SouthernBiotech, USA) and imaged on a laser scanning confocal microscope (A1, Nikon, Japan).

#### 2.4.6. Biopsy and immunoelectron microscopy

In addition to using immunohistochemical pathological method to identify NFTs described above, higher resolution immunoelectron microscopy was also employed to illustrate the ultrastructure of the identified NFTs. To get a biopsy sample for the test, on the day of sacrifice, after the SMI Monkey 10 was anesthetized with intramuscular injection of atropine (20 mg/kg, *i.m*.), ketamine (10 mg/kg, *i.m*.), and solium pentobarbital (25 mg/kg, *i.m*.), its head was fixed on a stereotaxic instrument and the skull on the frontal lobe was exposed under aseptic conditions by a middle line longitudinal skin incision followed by removal of the connective tissue. A small hole (2 mm in diameter) targeting the prefrontal cortex (Broadmann A9) was drilled on the skull according to the following stereotaxic coordinates, which had been obtained under MRI guidance (*35*): anterioposterior, +11.5 mm from the bregma; mediolateral, −11.3 mm from the sagittal sutur. A biopsy needle (14 G, TSK Corporation, Tochigi-ken, Japan) was inserted into the right frontal lobe at a depth of 12.5 mm to obtain biopsy samples of the prefrontal cortex, which were immediately transferred into phosphate buffer (PB, 0.1 M, pH7.4, containing 4% formaldehyde and 0.1% glutaraldehyde) and stored in the fixative for 12 h. The hole in the skull was filled with hemostatic sponge. Then the monkey continued to go through euthanasia and perfusion process.

The gray matter of the biopsy sample was cut into 0.1 mm^3^-small pieces and then post-fixed with 0.1% OsO_4_ for 30 min, at 4 °C. The fixed specimens were washed with 0.1 M PB and dehydrated in graded ethanol, infiltrated with 25%, 50% and 75% LR white acrylic resin (medium grade, Electron microscopy sciences, Cat. #14381-UC, diluted in ethanol) for 1 h, 2 h and 2 h, respectively. Then immersed in pure resin for 24 h and the resin was changed every 8 h. After that, the specimens were polymerized at 55 °C for 24 h at the bottom of closed gelatin capsules filled with fresh LR white resin.

Ultra-thin sections (90 nm) were cut by a Leica EM UC6 ultramicrotome (Leica, Shanghai, China), picked on 100 mesh nickel grids and processed for post-embedding immunogold labeling. The sections were incubated in PB buffer with 0.1 M glycine for 10 min to quench aldehydes, blocked with 5% BSA in PB buffer for 30 min, and then were incubated in the primary antibodies, AT8 (1:20), AT100 (1:100) or NFTs (1:100), diluted in PB buffer with 1% BSA overnight at 4 °C. After five washes with PB buffer and blocking with 1% BSA, the sections were incubated in secondary antibodies, goat-anti-mouse conjugate to 10 nm gold (1:40) or goat-anti-rabbit conjugate to 20 nm gold (1:40) diluted in PB with 1% BSA for 2 h, washed five times with PB buffer, and then fixed with 0.1% glutaraldehyde. After staining with uranyl acetate, the labeled grids were viewed using a JEOL 1230 (JEOL, Japan) at 120 KV. Under low magnification (30-60k), bundles of fibrous structures were localized first within the neuron’s somata and processes, and then the presence of gold nanoparticles was searched. Once the dense and well-defined black particles around 20 nm or 10 nm sizes were identified, they were further confirmed using a high-power field of vision (>80k). When the suspected particles appeared hollow ring under defocus, they were determined as gold nanoparticles. Images were collected using a Gatan 830 camera (Gatan, USA).

#### 2.4.7. Image analysis

The density of 6E10 stained SPs, TUNEL labeled cells and ratio of AT8 immunoreactive areas on analyzed visual fields in the Hp (hippocampus), EC (entorhinal cortex), TC (temporal cortex), PFC (prefrontal cortex) and OC (occipital cortex) (Figure S1) were quantitatively analyzed. In order to make the analysis accurate, the ROI (region of interest) of these five brain regions for statistical analysis was strictly defined based on the brain atlas (Paxinos G. et al. Rhesus monkey brain in stereotaxic co-ordinates. 1st ed. Academic Press; 1999). In the quantification of AT8 immunoreactive areas and TUNEL labeled cells, the ROI of Hp was defined as the CA4 region surrounded by dentate gyrus (*36*). In SPs counting, the whole hippocampus was selected as the ROI for the analysis. The ROI of EC was the grey matter region between the subiculum and rhinal fissure. The ROI of TC was the grey matter region of superior temporal gyrus, between the lateral fissure and superior temporal sulcus. That of the PFC was the grey matter region of dorsal lateral prefrontal cortex, between the superior arcuate sulcus and principal sulcus. And the ROI of the OC was the grey matter region of primary visual cortex, between the superior calcarine sulcus and external calcarine sulcus.

For 6E10 stained SPs counting, 10 sections (random coronal sections from the ROI of each monkey) were imaged with a 10× objective under the same acquisition parameters. Then the number of deposits stained by 6E10 of at least 10 μm in diameter in these 10 sections were counted manually and the density was calculated. Values derived from the 10 sections/region were averaged and presented as mean±SEM. Immunostaining was performed in parallel in slides from all animals.

AT8 immunoreactive areas were quantified by using particle analysis tool in the Image J (NIH software) (*36*). Briefly, first, 10 immunostained sections (random coronal sections from the ROI of each monkey) were imaged with a 20× objective under the same acquisition parameters. 50 fields (5 fields/section) of each ROI were selected randomly and converted into 8-bit gray scale and thresholds were adjusted and kept constant to highlight immunostained cells. Next, the immunoreactive particles were analyzed using the particle analysis tool. The ratio of the immunoreactive area over each analyzed visual field was calculated. The total area measured on each ROI was 1 mm^2^. Values derived from the 10 sections/region were averaged and presented as mean±SEM. Immunostaining was performed in parallel in slides from all animals.

For counting apoptotic cells, we used Image J (NIH software) to manually count the cells labeled with TUNEL. For every ROI of each monkey, 10 sections were randomly selected for staining, and images of 5 visual fields were randomly collected for each section. All the images were captured with a 20× objective under the same acquisition parameters. The total area measured on each ROI was 1 mm^2^. Values derived from 10 sections/region were averaged and expressed as mean±SEM. Labeling was performed in parallel in slides from all animals.

All the analyses were performed by an experimenter blind to animals’ individual information.

### 2.5. Statistics

All statistical analyses were carried out with the SPSS statistical software package (version 20, IBM, USA). The performance of the monkeys in the HIT, the density of SPs, the percentage of phosphorylated tau protein immunoreactivity areas and the number of apoptotic cells were analyzed by t-test. All the data were normally distributed (Kolmogorov-Smirnov: all *P* > 0.05). The alpha level was set at *P=*0.05 and all *P*-values were generated using two-tailed tests.

## 3. Results

To search naturally occurring AD monkeys, we first identified cognitively impaired aged monkeys with the tasks that assessed their visual-spatial working memory and emotional functions, which are two core AD clinical symptoms. Then, we carried out pathological tests to demonstrate those individuals with cognitive impairments were also associated with classical AD pathological hallmarks, including NFTs, SPs and neuronal loss. According to the diagnostic criteria of human AD, monkeys with all features mentioned above should be naturally occurring AD monkeys.

### 3.1. Three aged monkeys suffered from severe memory impairments (SMIs)

Memory impairments are the core clinical symptoms and the most important diagnostic criteria of AD. In this study, ten aged monkeys (26-29 years old) were selected to perform the VSDRT (Variable Spatial Delayed Response Task), a classical cognitive test that evaluates monkey’s spatial working memory by forcing the monkey to keep a piece of spatial information for a time period. The spatial working memory capacity of each monkey, which was measured by the maximum B value (the largest B value that each individual monkey was able to hold the spatial information at 90% correct or above for at least 5 consecutive days during the VSDRT testing stage, for details, see methods) is listed in Table 1. According the maximum B values, the 10 monkeys could be categorized into three groups: normal aged (the maximum B value between 35-38 seconds), mild memory impairment (MMI, the maximum B value between 14-16 seconds) and severe memory impairment (SMI, the maximum B value between 6-8 seconds) monkeys. The B value progress curves of the ten monkeys during the VSDRT testing stage are shown in Figure 1a. The ten curves again could be divided into same three groups as above: Monkey 1, 2, 3, 4 and 5 had normal working memory with the fastest progress curve and an average maximum B value of 36±0.63 seconds; Monkey 6 and 7 had MMI with middle progress curve and an average maximum B value of 15±1 seconds; Monkey 8, 9 and 10 had SMI with the slowest progress curve and an average maximum B value of 7±0.58 seconds.

**Figure 1.**
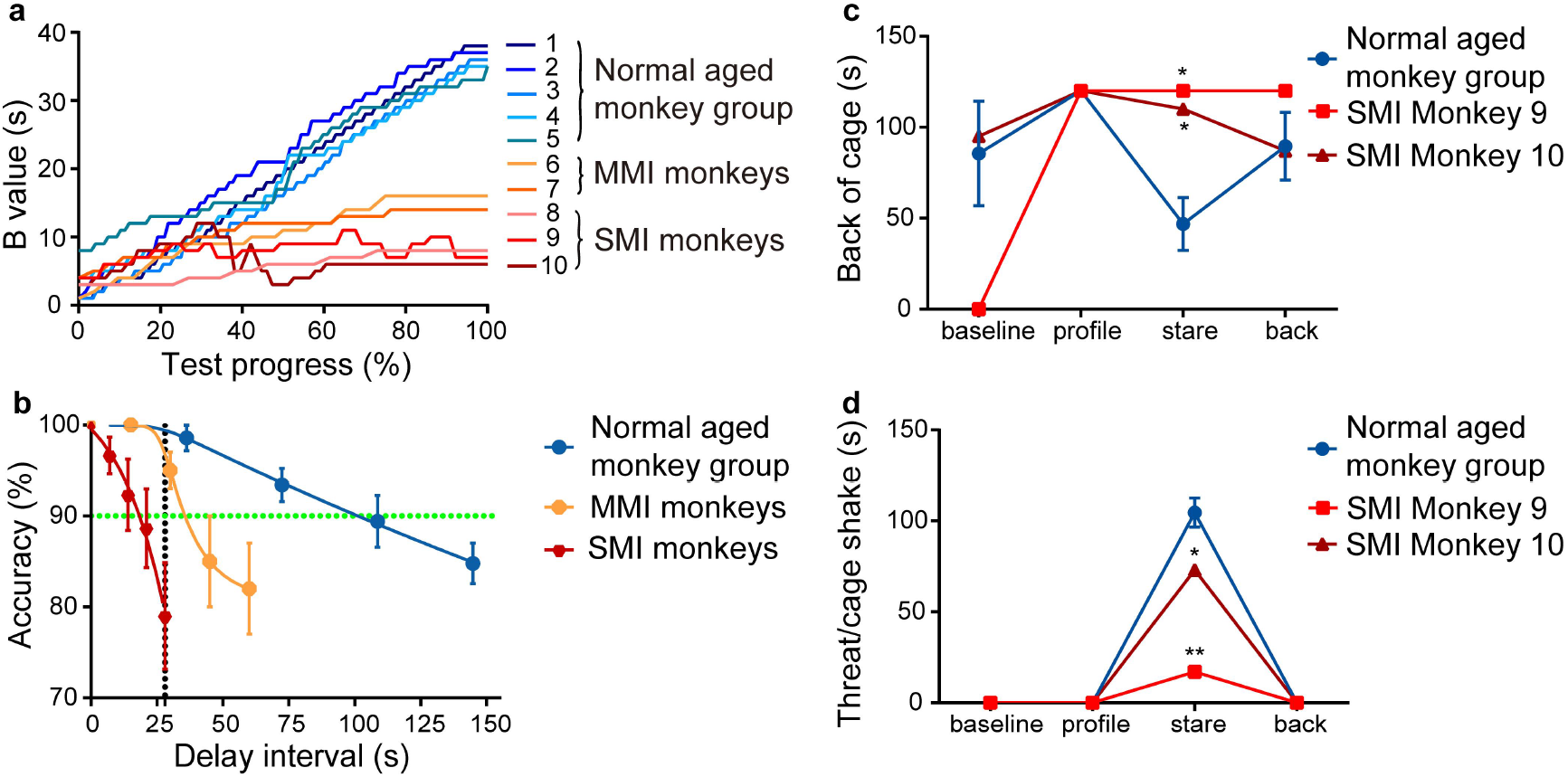
Performances of the aged monkeys in Variable Spatial Delay Response Task (VSDRT) and Human Intruder Test (HIT). (a) Progress curves of B value of the ten aged monkeys during the testing stage of the VSDRT. The monkeys could be clearly categorized into three groups according to both their maximum B values and the pattern of the curves. The Y-axis is the B value of each monkey. The average maximum B values of five normal aged monkeys was 36±0.63 seconds, the MMI monkeys’ maximum B values 15±1 seconds, and the SMI was 7±0.58 seconds. The X-axis is the progress of the testing stage, which is defined by experimental days of reaching certain B value/experimental days reached the maximum B value. Because each monkey achieved different B value at the end of the 1000-trial training stage, the initial B value of each monkey was different at the beginning of the testing stage (see methods). (b) The accuracy of the normal, MMI and SMI group at five different delay time periods (0×B; 1×B; 2×B; 3×B; 4×B) in the VSDRT when they had reached the maximum B values, respectively. The Y-axis is the performance accuracy in the VSDRT test. The X-axis represents the delay time periods in the task. The five dots of the blue curve show the accuracy and interval time at the different delay time period of the normal aged group, which are 0×B/ 100%/ 0 s, 1×B/ 98.6%/ 36.2 s, 2×B/ 93.4%/ 72.4 s, 3×B/ 89.4%/ 108.6 s, 4×B/ 84.8%/ 144.8 s, respectively. For the yellow curve of the MMI group are 0×B/ 100%/ 0 s, 1×B/ 100%/ 15 s, 2×B/ 95%/ 30 s, 3×B/ 85%/ 45 s, 4×B/ 82%/ 60 s, respectively. And for the red curve of the SMI groups are 0×B/ 100%/ 0 s, 1×B/ 96.67%/ 7 s, 2×B/ 92.33%/ 14 s, 3×B/ 88.67%/ 21 s, 4×B/ 79%/ 28 s. This figure shows that when the monkeys reached its longest delay (4×B), the normal aged group could maintain a longer delay period and higher accuracy than the MMI and SMI groups, and the MMI group could maintain a longer delay period and higher accuracy than the SMI group. The green dotted line indicates a 90% accuracy level. The black dotted line indicates the 4×B delay time period of the SMI group, which was 28 seconds. Note that the normal aged group could reach the 100% correct at this delay time period, while the MMI monkeys and SMI monkeys could only reach 96% and 80% correct rate, respectively. (c-d) In the Human Intruder Test (HIT), compared with the normal aged monkey group (n=4), the SMI Monkey 9 and 10 spent more time on staying at back of the cage (Monkey 9, *P=*0.015; Monkey 10, *P=*0.023; one-sample t-test) and spent less time on threating/cage shaking (Monkey 9, *P=*0.002; Monkey 10, *P=*0.03; one-sample t-test) in the stare phase, suggesting that the SMI Monkey 9 and 10 were significantly less responsive to the stressful stimuli. There were no significant differences in the profile and back phases between the normal and SMI monkeys. The Y-axis is the total time of each behavior displayed by the monkeys. The X-axis is the different stages of the HIT. All data are mean±SEM. *: *P*<0.05.

In addition to the differences in maximum B values and progress curves, the normal aged monkeys were also able to maintain a higher successful rate during the longer delay periods than the memory impaired ones. As showed by the green dotted line in Figure 1b, the normal aged monkeys were able to maintain a 90% correct rate when the delay time was 100 seconds. But for the same 90% correct rate, the MMI monkeys could only last 35 seconds, while the SMI monkeys about 20 seconds. Also, in Figure 1b, the black dotted line shows that, for the delay time that the normal aged monkeys were able to perform 100% correct rate, the MMI monkeys and SMI monkeys could only reach 96% and 80% correct rate, respectively. Another noteworthy fact of this figure is that the correct response rates of all animals at a 0-second delay time period are 100%, indicating that, except for the memory impairment, the visual and motor functions of all the 10 aged monkeys are normal. Taken together, the data suggest that Monkey 8, 9 and 10 of the SMI group have the highest chance to have AD.

### 3.2. the SMI monkeys also displayed apathy

Since decreased responsiveness to environmental changes (*apathy*) is the most common behavioral disorder next to memory impairments in AD patients (*4*), the Human Intruder Test (HIT) was employed in this study to quantify the monkey’s response to environmental stimuli. The HIT is a widely used behavioral test to assess monkeys’ reaction to a sudden change of the environment by exposing it to an unfamiliar person (intruder). A normal monkey tends to show some anxious behaviors to the stare of the intruder as manifested by its increased threatening behavior (*34*). We wanted to investigate whether the SMI monkeys responded differently to an intruder than the normal aged monkeys. The normal aged Monkey 1, 2, 3 and 5 and the SMI Monkey 9 and 10 were involved in this test. For the normal aged Monkey 4 and the SMI Monkey 8 (two of the oldest individuals in the group) became very weak at the time of the HIT test. To ensure the data quality, they were excluded from the test.

This test took place after the VSDRT had been completed. After initial data sorting, only two kinds of behaviors: threating/cage shaking and staying at the back of cage, which were the dominant reactions and displayed by all of the 6 tested monkeys were compared between the SMI and control monkeys. The results showed that the two groups responded differently to the staring of the intruder (Figure 1c and 1d). Compared with the 4 normal aged monkeys as a group, the SMI Monkey 9 and 10 were less responsive to the staring intruder as demonstrated by spending more time just staying at back of the cage (Figure 1c, *P=*0.023, one-sample t-test), and spent less time in threating/cage shaking (Figure 1d, *P=*0.03, one-sample t-test) during the stare phase. The staying at the back of its cage represented the monkey’s mild reaction (a low reaction) to the intruder (*33*), and threating/cage shaking is an exaggerated aggressive response to the intruder, representing a high reaction (*34*). These two behaviors antagonize each other because they were the dominant reactions among the tested monkeys. While staying at the back of its cage increased, the threating/cage shaking decreased, indicating the animals were insensitive to the environmental stimuli. Further video analysis showed that the SMI Monkey 9 and 10 had not manifested any fear behaviors, such as lipsmack, teeth gnash and fear grimace, during the stare phase. Thus, no responding to the staring, instead of fearing of it, was the most plausible explanation of the differences between the two SMI monkeys and the controls (see Movie 1 in Supplementary Materials).

On the other hand, there were no significant differences in the profile and back phases between the normal aged group and the SMI monkeys, suggesting that the SMI Monkey 9 and 10 responded to a non-stressful environmental stimulus the same way as the controls. Taken together, these results suggested that the two SMI monkeys were significantly less responsive to stressful environmental stimuli, showing some apathy, which, in addition to memory deficits, is also a typical clinical symptom of AD patients (*4*).

### 3.3. All the SMI monkeys had large amount of neurofibrillary tangles in their brains

The NFTs, which are made up of hyperphosphorylated tau proteins, are one of the two most important pathological hallmarks of AD. To determine the presence of tau pathology in the brain of the SMI monkeys, we stained brain sections of the SMI and normal aged monkeys with monoclonal antibody AT8, a widely used antibody for hyperphosphorylated tau protein detection and were used in several tau-pathological studies in Macaca monkeys (*37-40*), which recognizes phosphorylation of tau at serine 202/threonine 205, to illustrate the presence of hyperphosphorylated tau.

The results revealed that all the SMI monkeys had a large number of hyperphosphorylated tau proteins in AD-related brain regions, including the Hp, EC, TC and PFC. As an example, Figure 2a and 2b show the AT8 staining of the TC of one SMI monkey (Monkey 8) and one normal aged monkey (Monkey 4). The SMI Monkey 8 had much more AT8 positive cells than the normal aged Monkey 4 did in this area. Further quantitative analysis in Figure 2c shows that the percentage of AT8 positive area of the SMI Monkey 8 in the TC is significantly higher than that of normal aged monkey group (n=3) (*P*=0.01, one-sample t-test). To illustrated group data, Figure S2 shows the AT8 staining of the AD related region Hp, EC, TC and PFC of all the 3 SMI monkeys and the 3 normal aged monkeys, as well as quantitative analyses of the percentage of AT8 positive areas in each of the brain regions mentioned above. In Figure S2a, the numbers of AT8 positive cells in the Hp, EC, TC and PFC of the SMI monkeys are much larger than those of the normal aged monkey group. The quantitative results show that except for the TC of Monkey 9, the percentages of AT8 positive area in the Hp, EC, TC and PFC of the SMI monkeys were higher than those of the normal group. Among which, the AT8 positive areas in the Hp, TC and PFC of SMI Monkey 8 and 10 and the EC and PFC of Monkey 9 were statistically significant (Figure S2b-d).

**Figure 2.**
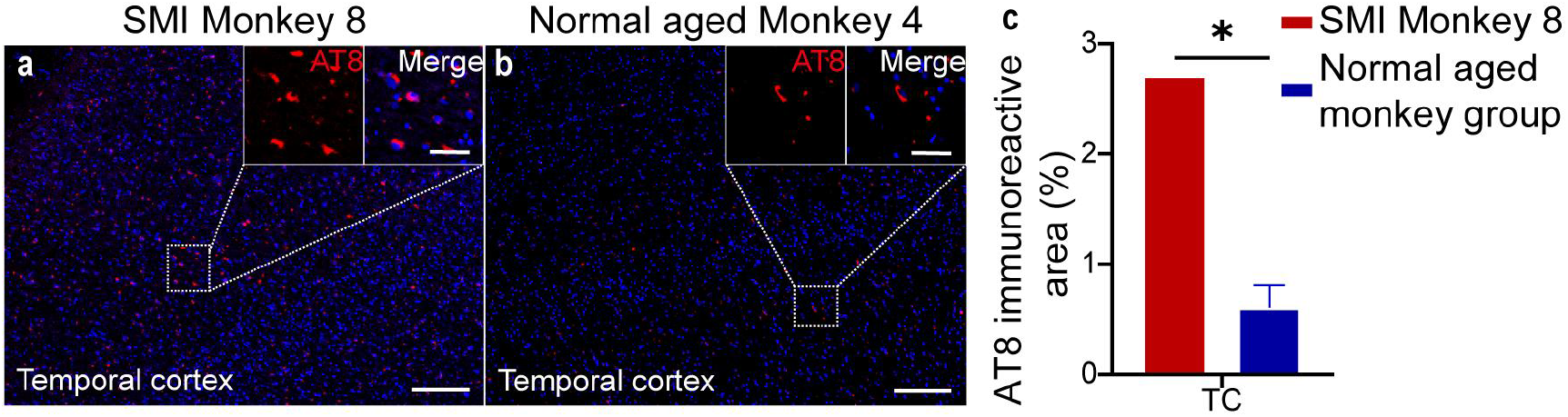
AT8 immunofluorescence staining images of the temporal cortex (TC) of the SMI Monkey 8 and normal aged Monkey 4 and a quantitative analysis of AT8 immuno-positive area in the TC between the SMI monkey 8 and the normal aged monkey group (n=3). (a, b) Representative images of hyperphosphorylated tau proteins (red, AT8 monoclonal antibody) in the TC of the SMI Monkey 8 and normal aged Monkey 4, showing that the amount of AT8 positive cells in the TC of the SMI Monkey 8 is significantly larger than that of normal aged Monkey 4. Scale bars: 200 μm in general view; 50 μm in larger view. (c) Comparison of the percentage of the AT8 immuno-positive area in the TC between the SMI Monkey 8 and the normal aged monkey group (n=3), showing that the percentage of AT8 positive area of the SMI Monkey 8 in the TC is significantly higher than that of normal aged group (*P*=0.01; one-sample t-test; for details, please see Methods). The data of normal aged monkeys is mean±SEM. *: *P*<0.05.

In addition to above immunohistochemical demonstration of NFTs, higher resolution immunoelectron microscopy was also employed to illustrate the ultrastructure of NFTs identified in the PFC of the SMI Monkey 10. The immunoelectron microscopy was performed using primary antibodies of NFTs (anti-neurofibrillary tangles polyclonal antibody, Figure 3a), AT100 (Figure 3b, recognize phosphorylation of tau at serine 212 and threonine 214) and AT8 (Figure 3c, recognize phosphorylation of tau at serine 202/ threonine 205) and secondary antibodies conjugated to 10/20 nm gold to illustrate the NFTs (*41, 42*). Under low magnification (30-60 k), bundles of fibrous structures were localized in the neurons’ soma and processes, and the presence of gold nanoparticles was searched. Once dense and well-defined black particles around 20 nm or 10 nm sizes were found, they were further confirmed using a high-power field of vision (>80 k). When the candidate particles appeared hollow ring under defocus, they were determined as nano-gold particles. The antibody-conjugated nano-gold particles (red arrows in Figure 3 a-c) marking the NFTs (by NFTs antibody) or hyperphosphorylated tau proteins (by AT100 and AT8 antibodies) were observed within tangle-like structures in the neuronal soma, illustrating the ultrastructure of the NFTs in the PFC of the SMI Monkey 10. These ultra-structures observed here are similar to the tangle structures demonstrated previously in AD patient brains and aged macaques using similar technology (*19, 43*). Therefore, we not only found region-specific NFTs in SMI monkeys, but also illuminated the submicroscopic structure of the NFTs, which further proved that the SMI monkeys had NTFs in the brain.

**Figure 3.**
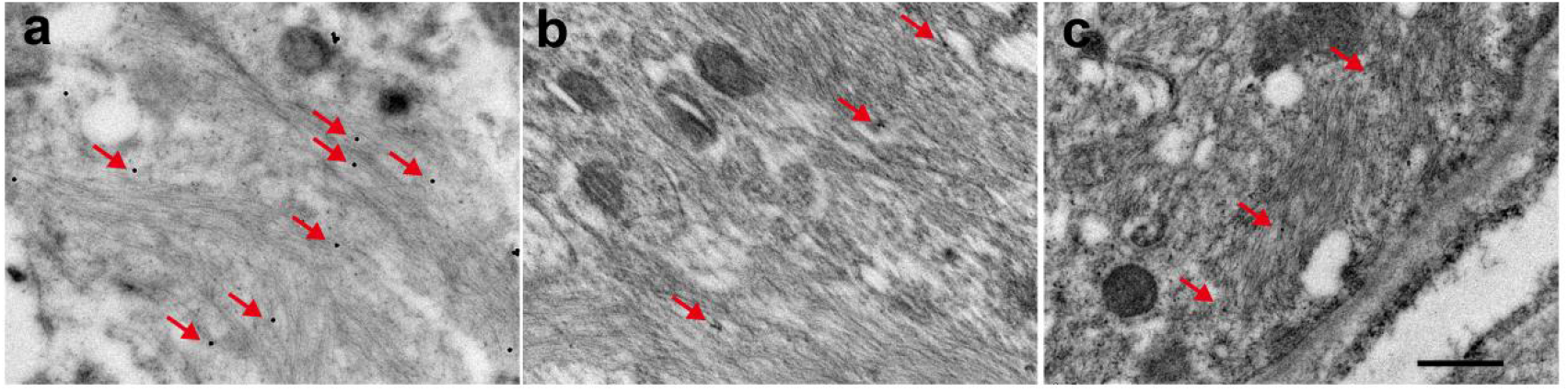
The ultrastructure of NFTs in the PFC of the SMI Monkey 10 illustrated by immunoelectron microscopy. Immunoelectron microscopy was performed with antibodies NFTs (a), AT100 (b) and AT8 (c), which recognized the neurofibrillary tangles, hyperphosphorylated tau in Ser 214/Thr 212 and Ser202/Thr205, respectively. The antibodies were conjugated to 10 (b and c) or 20 (a) nm gold which can be detected as dense and well-defined black particles under electron microscopy. The red arrows in a-c mark the nano-gold particles in the tangle-like structures within neuronal soma, showing the ultrastructure of the NFTs. Scale bar: 0.5 μm.

### 3.4. Some of the SMI monkeys had abundant Aβ plaques in their brains

As the SPs, consisted of amyloid deposition, are the other most important pathological feature of AD, we also investigated them in the pathological assessment. Since the cerebral cortex, in particular the isocortex, is the predilection site for the deposition of SPs (*44*), the pathological assessment of SPs was carried out on the Hp, EC, TC and PFC. The monoclonal antibody 6E10, one of the first cloned and commercialized monoclonal antibodies against Aβ and used commonly in Aβ-pathological studies of Macaca monkeys (*38, 39, 45*) which recognizes Aβ 1-16 sequence of amino acides, was used to stain the SPs in this study.

The results demonstrated that SPs pathology was also presented in the brain of two out of the three SMI monkeys. As an example, Figure 4a and b showed the 6E10 staining images of the TC of one SMI monkey (Monkey 8) and one normal aged monkey (Monkey 4). The SMI Monkey 8 had a large number of SPs in the TC (Figure 4a), whose size and number were similar to those typical SPs in AD patients(*44*). These SPs included dense plaques and diffused plaques in brain parenchyma or around blood vessels (Figure 4c-f). No 6E10 staining SPs was found in the TC region of the normal aged Monkey 4 (Figure 4b). The quantitative analysis in Figure 4g showed that the density of SPs in the TC of the SMI Monkey 8 was significantly higher than that of normal aged monkey group (n=3, one-sample t-test). Studies have shown that SPs often recruit glial cells and appear as localized inflammatory responses in AD patients by using immunofluorescence co-stain the SPs and gliocyte to visualize this local inflammatory response (*46*). In the TC of the SMI Monkey 8, prominent aggregation of glial cells could be seen around the SPs by GFAP (recognizing astrocyte), Iba1 (recognizing activated microglia) and 6E10 immunofluorescence co-staining. Figure 4h and 4i show the reactive-phenotype astrocytes with hypertrophied processes (green, marked by white arrow) envelop and penetrated into the plaques (6E10, red). Figure 4j and 4k show the activated microglia (green, marked by white arrow) gathered near the plaques (6E10, red). Notably, some Aβ was inside these microglia (the white arrowheads in Figure 4k), suggesting that the activated microglia might be phagocytizing the plaques. As group data, Figure S3 shows 6E10 staining images of the AD related cortical regions of Hp, EC, TC and PFC of all the three SMI monkeys and the three normal aged monkeys, as well as the quantified SPs’ density in each of the brain regions of the 3 SMI monkeys compared with that of the 3 normal aged monkeys as a group (one-sample t-test), respectively. In Figure S3a and b, the number of SPs in TC and PFC of the SMI Monkey 8 was significantly higher than that of normal aged monkeys. On the other hand, the SMI Monkey 10 had a mild SPs pathology in the TC and PFC (Figure S3a and S3d), and there were no detectable SPs in the SMI Monkey 9 (Figure S3a and S3c). In the normal aged group, the amyloid deposition was absent in normal Monkey 1 and 4, and a small number of SPs was observed in the TC and PFC of normal Monkey 2 (Figure S3a). This is consistent with data from AD patients. Some autopsy studies report that some aged people have SPs or/and NFTs pathology in their brains with normal cognition (*21*). In summary, the SPs’ study here revealed that two out of the three SMI monkeys (Monkey 8 and 10) showed clear SPs pathology in the AD related brain regions.

**Figure 4.**
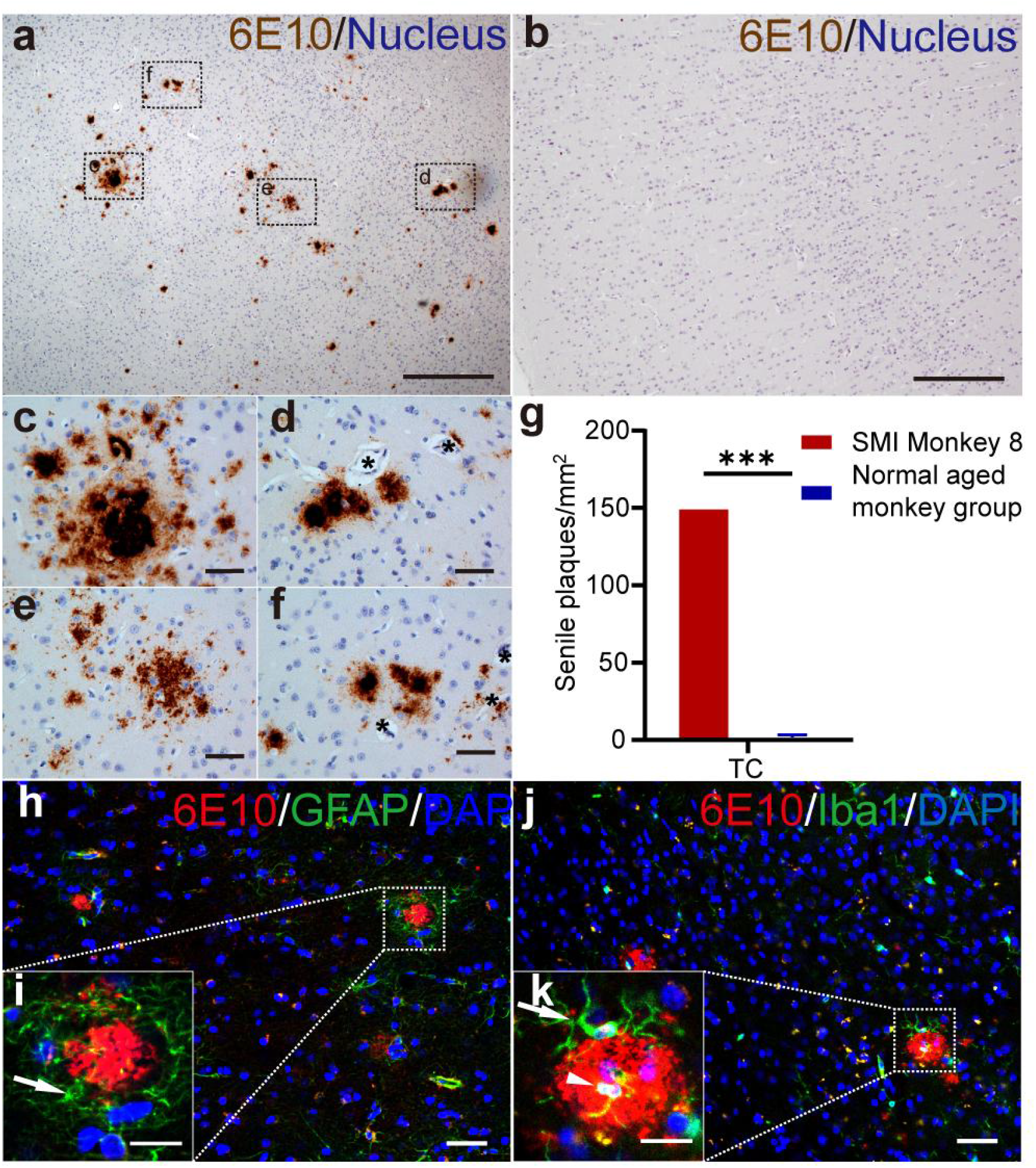
6E10 staining images of the temporal cortex (TC) SPs of the SMI Monkey 8 and normal aged Monkey 4 and a quantitative SPs analysis between the SMI Monkey 8 and normal aged monkey group (n=3). (a, b) Representative images of SPs (brown, 6E10 monoclonal antibody) form the TC of the SMI Monkey 8 and normal aged Monkey 4, show that there are a large amount of 6E10 stained SPs in the TC of the SMI Monkey 8 and the absence of SPs in the same region of the normal aged Monkey 4. Scale bars: 500 μm. (c-f) Enlarged views of SPs in the dot-lined squares in (a) illustrate dense plaques (c, d and f) and diffused plaques (e) in brain parenchyma (c, e and f) or around blood vessels (d and f), respectively. Stars in (d and f) mark the blood vessels surrounded by SPs. Scale bars: 50 μm. (g) Comparison of the SPs’ density in the TC between the SMI Monkey 8 and normal aged monkey group (n=3), showing that the SPs’ density of the SMI Monkey 8 in the TC is significantly higher than that of normal aged group (*P* < 0.001; one-sample t-test, for details, please see Methods. The data of normal aged monkey group is represented as the mean±SEM. ***: *P*<0.001. (h-k) Confocal micro-images of the TC of the SMI Monkey 8 show co-localization of 6E10 (red), GFAP (green in h and i) and Iba1 (green in j and k), showing the SPs surrounded by activated glial cells (marked by white arrows). Especially, white arrowhead in (k) indicates the co-stain of 6E10 and Iba1, suggesting the phagocytizing of the SPs by activated microglia. Scale bars: 200 μm in (h and j); 20 μm in (i and k).

### 3.5. All the SMI monkeys had severe cell apoptosis

Brain atrophy is another important pathological feature of AD, and to detect brain cell apoptosis is a direct way to illustrate brain atrophy (*47*). Many cells in AD patients have been shown to exhibited terminal deoxynucleotidyl transferase (TdT) labeling for DNA strand breaks, which reflects neuronal vulnerability and cell loss associated with brain atrophy (*48*). We compared the TdT labeling in the AD related Hp, EC, TC and PFC cortical regions of all the 3 SMI monkeys and 3 normal aged monkeys using Terminal-deoxynucleotidyl Transferase Mediated Nick End Labeling (TUNEL).

Abundant TdT labeled cells were identified in the Hp, EC, TC and PFC of the 3 SMI monkeys. As an example, Figure 5a and 5b show the TdT labeled images from the TC of the SMI Monkey 8 and normal aged Monkey 4. The number of TdT labeled cells in the SMI Monkey 8 is much larger than that of the normal aged Monkey 4. Quantitative results in Figure 5c reveals that the density of apoptotic cells in the TC of the SMI Monkey 8 is significantly higher than that of normal aged group (one-sample t-test). To illustrated group data, Figure S4 shows the TdT labeling of the AD related cortical region Hp, EC, TC and PFC of all the 3 SMI monkeys and 3 normal aged monkeys, and the quantitative analyses of the TdT-labeled-cell’ density in each of the brain regions. In Figure S4a, the number of TdT labeled cells in the Hp, EC, TC and PFC of the three SMI monkeys are much higher than that in the normal aged group. Quantitative results showed that except for the Hp and PFC of SMI Monkey 10, the densities of TdT labeled cells in the Hp, EC and TC and PFC of the SMI monkeys are significantly higher than the average of the 3 normal aged monkeys as a group (Figure S4b-d).

**Figure 5.**
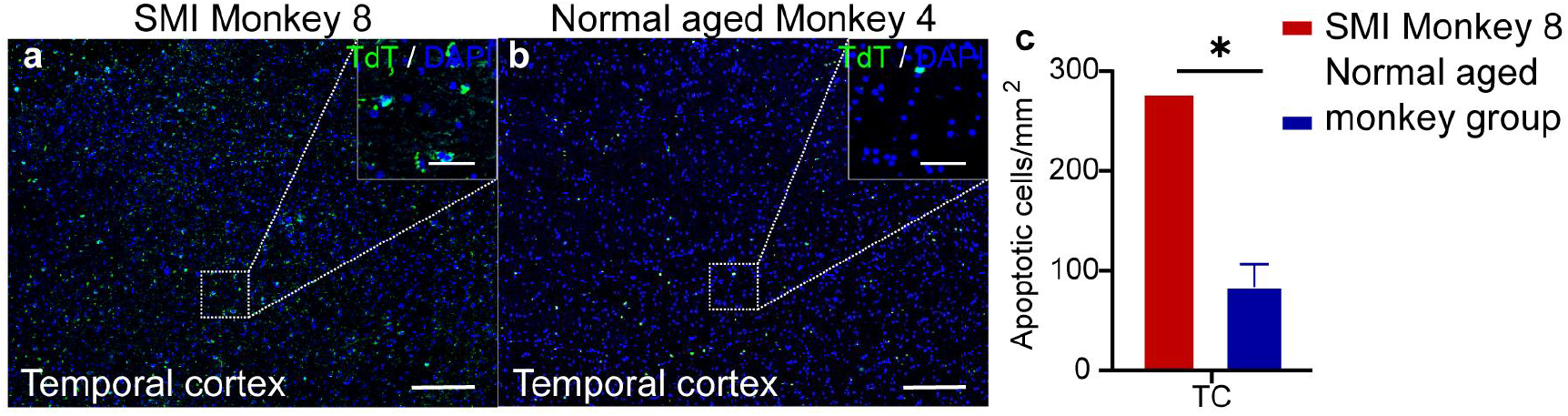
TdT labeling of apoptotic cells in the TC of the SMI Monkey 8 and normal aged Monkey 4 and related quantitative analysis between the SMI Monkey 8 and normal aged monkey group (n=3). (a, b) Representative images of TdT labeled apoptotic cells (green) in the TC of the SMI Monkey 8 and normal aged Monkey 4, showing that the amount of TdT labeled apoptotic cells in the TC of the SMI Monkey 8 is significantly larger than those in normal aged Monkey 4. Scale bars: 200 μm in general view; 50 μm in larger view. (c) Comparison of the apoptotic cells’ density in the TC between the SMI Monkey 8 and the normal aged monkey group (n=3), showing that the density of the SMI Monkey 8 in TC is significantly higher than that of normal aged group (*P*=0.014, one-sample t-test, for details, please see Methods). The data of normal aged monkeys is mean±SEM. *: *P* < 0.05.

### 3.6. The two SMI monkeys that had all the AD core hallmarks (both the clinic symptoms and pathologic changes) also had similar cortical regional pathological distribution patterns as AD patients

After the pathological studies above, it was clear that two (SMI Monkey 8 and 10) out of the three SMI monkeys had all the three AD core pathologic hallmarks, namely NFTs, apoptosis and SPs (Figure 2-5 and S2-4). According to the diagnostic criteria of AD (*22, 23*), these two aged rhesus monkeys were clearly spontaneous AD cases. In AD patients, the progress of tau pathology and apoptosis are positively correlated with the cognitive impairment, and the brain regions such as the Hp, EC, TC and PFC are more susceptible to the affliction of the tau pathology and apoptosis than visual cortical area OC (occipital cortex) (*44, 48*). Thus, the distributions of hyperphosphorylated tau proteins and TdT labeled cells in the AD related (Hp, EC, TC and PFC) and less-related brain region (OC) of these two AD monkeys are analyzed quantitatively and it was found that their distributions were consistent with the patterns of AD patients. That is, the visual area OC was less vulnerable than the other cognitive related regions (Figure 6).

**Figure 6.**
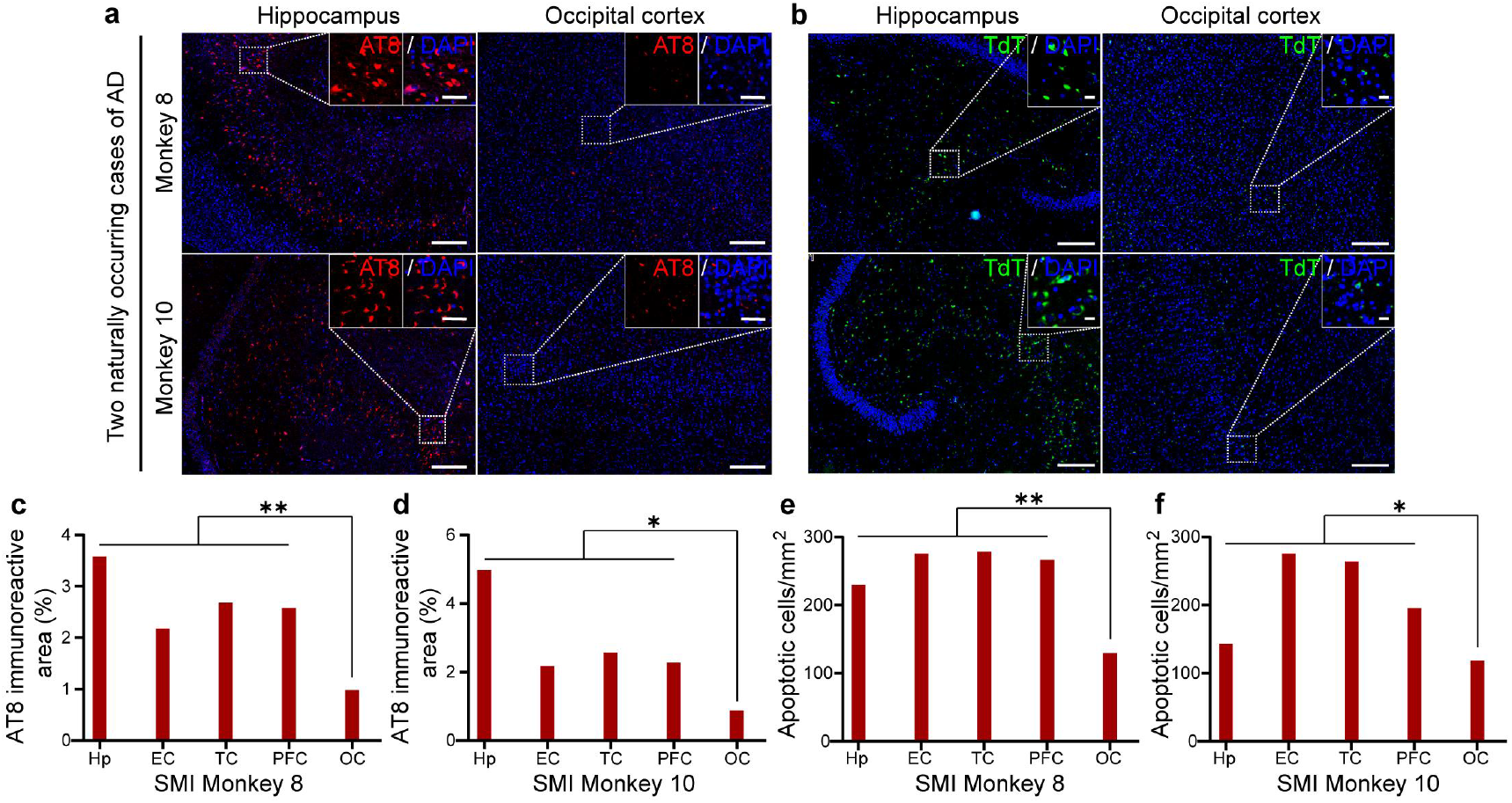
AT8 immunofluorescence staining and TdT labeling of apoptotic cells of the AD related region Hp and AD unrelated region OC of the SMI Monkey 8 and 10 and their quantitative analyses between the AD related regions Hp, EC, TC and PFC and unrelated region OC. Representative images of AT8 immunofluorescence staining (red) and TdT labeled apoptotic cells (green) in the Hp and OC of the SMI Monkey 8 and 10, showing that the numbers of AT8 positive or TdT labeled apoptotic cells in the Hp are significantly larger than those in the OC. Scale bars: 200 μm in general view; 50 μm in larger view. (c-f) Quantitative analyses of the percentage of the AT8 immuno-positive area and the apoptotic cell density in the Hp, EC, TC, PFC and OC of the SMI Monkey 8 and 10, respectively, showing that the AT8 positive area/ apoptotic cell density in the AD related region Hp, EC, TC and PFC are significantly higher than these of the AD unrelated region OC (in c *P*=0.009, in d *P*=0.049, in e *P*=0.001 and in f *P*=0.047, one-sample t-test, for details, please see Methods).

Figure 6a and 6b show the AT8 staining (hyperphosphorylated tau proteins) and TdT labeling (apoptosis) images in AD related region Hp and unrelated region OC of the SMI Monkey 8 and 10. The SMI Monkey 8 and 10 have much more AT8 positive cells and TdT labeled cells in the Hp than in OC. Further quantitative analyses in Figure 6c-f show that the percentage of AT8 positive area and the density of TdT labeled cells in the AD related regions Hp, EC, TC and PFC of SMI Monkey 8 and 10 are much higher than those in AD unrelated region OC. After comparing the OC with AD related regions (Hp, EC, TC and PFC as a group) by one-sample t-test, it was revealed that the percentage of AT8 positive area and density of TdT labeled cells in AD related regions were significantly higher than those of the OC in both SMI Monkey 8 and 10. These results suggest that in these two AD monkeys, similar to AD patients, the OC is less vulnerable to AD pathological changes than the Hp, EC, TC and PFC (*44, 48*).

## 4. Discussion

In our efforts to identify monkeys with naturally developed AD, we have acquired all the behavioral and pathological data in a subset of rhesus monkeys with advanced age that are essential and required for the AD classification presently in use of human medicine. In addition to an assessment of memory loss and other cognitive deficits such as apathy, we have also analyzed for the presence of critical pathological AD markers such as NFTs, SPs and neuronal loss. We have confirmed that two of the 10 aged monkeys (the SMI Monkey 8 and 10) in our study had typical AD clinical symptoms, including memory deterioration and apathy, as well as AD pathological hallmarks such as intracellular NFTs, extracellular SPs and significant neuronal loss. Furthermore, the distribution patterns of the NFTs and cell loss in these two SMI monkeys were consistent with the patterns of AD patients. That is, the visual cortical area OC was less vulnerable than other cognitive related areas. With all these findings, we were able to demonstrate, for the first time, that monkeys upon aging may develop spontaneous AD just like human beings.

Clinical trials with novel AD drug candidates have been mostly unsuccessful (*7, 9, 11*), which has been in part attributed to an over-reliance on inadequate rodent animal models (*49*). While there is 83% genetic similarity between mouse and human, there are significant differences in the CNS between the two species with only a 10% homology for co-expressed genes (*50*). It is therefore not surprising that the phylogenetic differences between rodents and humans have made the translation between the two species during drug development trials difficult and are likely responsible for the observed extremely high failure rates (99.6%) (*10*). Another important fact is that rodents do not develop AD naturally, but in all studies is artificially induced (*51*). Hence, from the perspective of translational medicine and drug development efforts, rodents are not the ideal species to serve as a model for AD (*52, 53*).

As a consequence of evolution, the high similarity between the human and monkey brains in terms of overall architecture and functional networking makes the monkey the preferred animal model for the study of AD progression and drug development (*54-56*). Aged monkeys, like aged humans, develop behavioral and cellular abnormalities over time that resemble AD (*56*). Although cognitive impairment and pathological hallmarks have been previously reported in separate aged monkeys (*18, 19, 57-59*), the scientific community has so far not accepted the fact that monkeys can suffer from AD naturally. It has been suggested that an improved understanding why AD fails to develop in species that are biologically proximal to humans, such as monkeys, could disclose new therapeutic targets involved neurodegeneration and dementia (*60, 61*). The most important reason for this skepticism is the fact that the four criteria that define AD, namely cognitive impairment, SPs, NFTs and neuronal loss, have never been demonstrated to exist in the same monkey. To address this issue, we have for the first time repeated the discovery process of human AD in monkeys and carried out a comprehensive investigation of AD hallmarks in the same cohort of aged monkeys. This investigation was conducted in a manner similar to the current clinical way to define AD by typical clinical symptoms combined with classical pathological markers (*22, 23*). Our data suggest for the first time that aged monkeys indeed could have NFTs, SPs and neuronal loss in their brain tissues, which were accompanied with core AD cognitive impairments. These results strongly suggest the existence of naturally occurring AD monkeys.

With regard to the cognitive impairments, we assessed with VSDRT the core clinical diagnostic symptom of AD patients, working memory capacities, which provided a classic indication on the monkeys’ visuospatial working memory ability (*30, 62-66*). Having demonstrated that three of the 10 aged monkeys showed severe memory impairment, we next focused on another behavioral abnormality in the AD patients, apathy behavior. Using HIT, a noninvasive tool widely used for assessing a monkey’s response to environmental stimuli (*32*), two of the above SMI monkeys that was available took the HIT and showed apathy-like behaviors such as lowered responsiveness during the stranger stare phase. Collectively, the SMI monkeys showed the classic cognitive impairments similar to those observed in AD patients.

For the pathological investigation, we focused on the Hp, EC, TC and PFC that are crucial for working memory function and AD development (*67, 68*) and found great numbers of both diffused SPs and dense SPs in the brain of two SMI monkeys, something that has been previously reported in aged monkeys (*15, 16, 18*). Significant NFTs were also present in all the three SMI monkeys’ brain as shown by immunostaining and immunoelectron microscopy. Such NFTs have only been reported in one previous study with macaques (*19*). Also, in the SMI monkeys, we found that apoptotic cells were prominent in the AD-associated brain regions. This important feature of neurodegenerative diseases has not been reported in aged monkeys before. Furthermore, the distribution patterns of the hyperphosphorylated tau proteins and apoptotic cells in these two AD monkeys were consistent with the patterns of AD patients. That is, the visual area OC is less vulnerable than other cognitive related areas (*44, 48*). These pathological studies suggested that two out of the three SMI monkeys had all the AD core features, namely NFTs, SPs and cell loss, which could be defined as naturally occurring cases of AD.

In summary, we have been able to show for the first time in aged rhesus monkey a clear convergence of core AD hallmarks that include cognitive impairments, NFTs, SPs and neuronal loss. These AD hallmarks were collectively and consistently presented in the same animal which justifies the claim that we have discovered for the first time two naturally occurring cases of AD in rhesus monkey. Based on the results we submit AD is not a uniquely human disease and provides us with the opportunity to study its pathogenesis and pathobiology from an evolutionary perspective. Comparative analyses on differences in AD phenotypes among primates will be of great benefit for AD studies, ultimately resulting in an improved understanding of disease development, treatments and prevention. Our results demonstrate that monkeys represent valid and reliable AD models that can fully mimic both abnormal behaviors and pathology observed in AD patients. They also have great potential for translational studies of AD as well as understanding the etiology and identification for early diagnostic biomarkers.

## COMPETING INTERESTS

The authors declare that they have no competing interests.

## AUTHORS’ CONTRIBUTIONS

Z.H.L., X.P.H., S.H.W., R.Y.H., H.L., Z.B.W. and L.M.W. performed the experiments. Z.H.L., X.P.H., S.H.W., R.Y.H., L.M.W., X.M. and D.D.Q. analyzed the data. Z.H.L., Y.K., Y.Q.G. and W.C.W. performed the immunoelectron microscopy. C.W.T., Z.Q.X., W.C.W. and X.T.H. designed the project. Z.H.L., C.W.T. and W.C.W. wrote the manuscript. Z.H.L., H.L., C.W.T., W.C.W. and X.T.H. revised the manuscript. All authors read and approved the final version of the manuscript.

## ACKNOWLEDGMENTS

We would like to thank technician Fang Gao, Xu Wang, Lijun Pan, Rong Sun and Chunying Yin for their help for data collection. We also thank Zhengfei Hu of Kunming Institute of Zoology, Chinese Academy of Sciences, for his kind help for animal care.

This study was supported partially by the National Science and Technology Innovation 2030 Major Program (2021ZD0200900), National Key Research and Development Program of China (2021YFF0702700, 2018YFA0801403), the Key-Area Research and Development Program of Guangdong Province (2019B030335001), National Natural Science Foundation of China (81941014, 81771387, 31800901, 31960178), Strategic Priority Research Program of the Chinese Academy of Sciences (CAS) (XDB32060200), Applied Basic Research Programs of Science and Technology Commission Foundation of Yunnan Province (202001AT070130, 202101AT070437, 202101AT070403), and CAS “Light of West China” Program.

## Supplementary Materials

**Figure S1.**
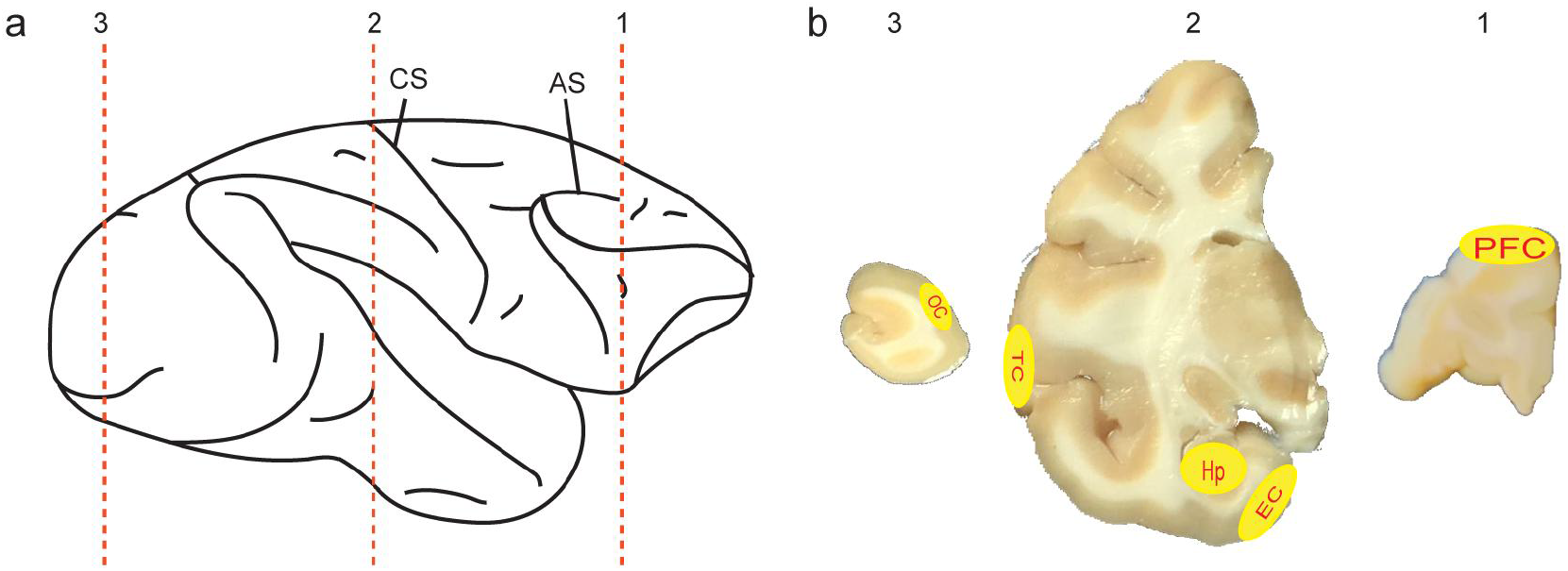
Schematic diagram illustrates the brain regions selected for the pathological tests. (a) Lateral view of the right hemisphere of a monkey brain. Red dashed lines show the locations from where brain slices were obtained for the pathological tests. AS: arcuate sulcus, CS: central sulcus. (b) The corresponding coronal sections from the dashed lines in (a). The areas marked yellow in each section were selected for the pathological tests. EC: entorhinal cortex, PFC: prefrontal cortex, TC: temporal cortex, Hp: hippocampus, OC: occipital cortex.

**Figure S2.**
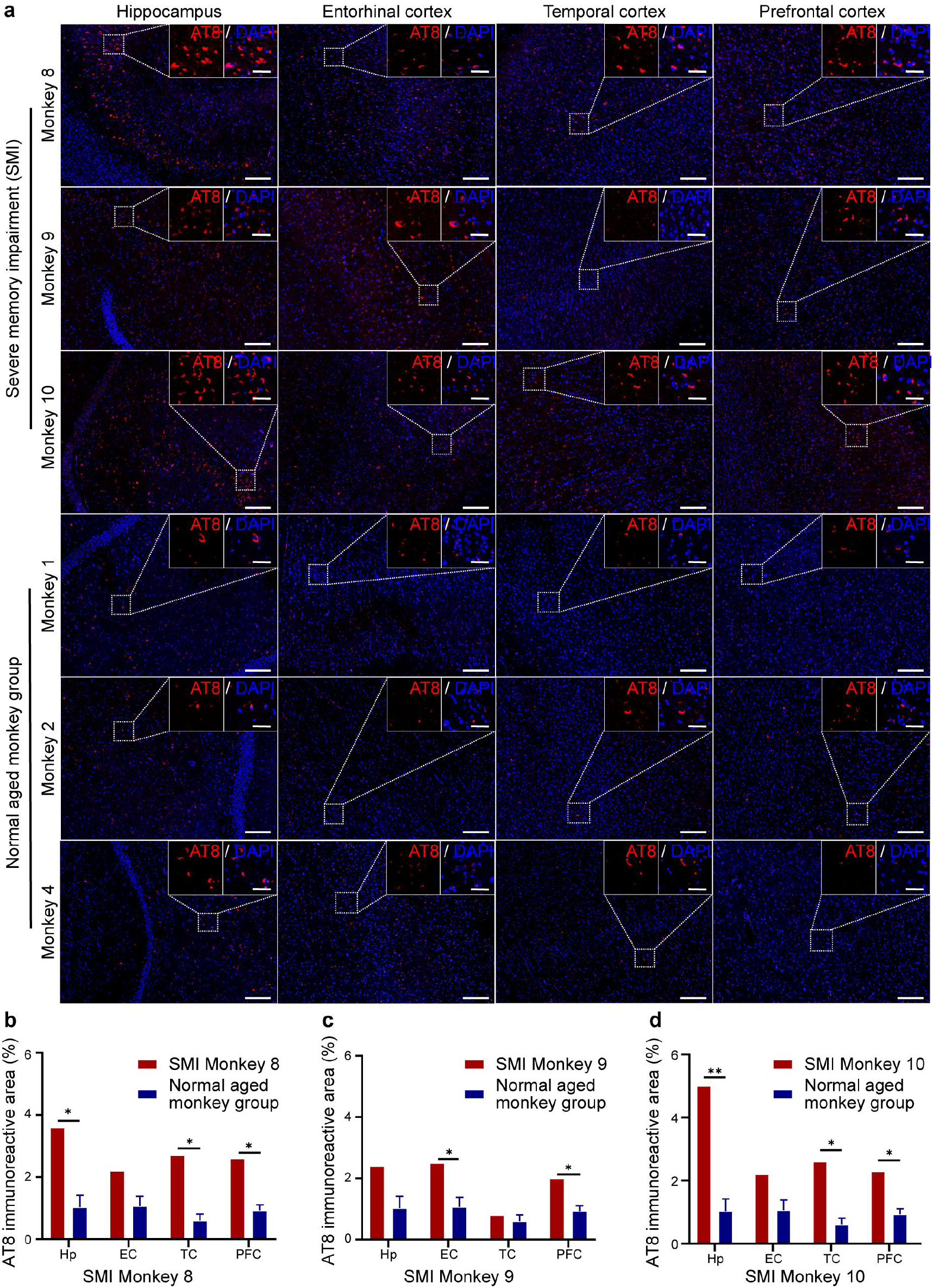
Immunofluorescence staining images and quantitative analyses of AT8 immuno-positive area in the AD related brain regions of the three SMI monkeys and the three normal aged monkeys. (a) Representative images of hyperphosphorylated tau proteins (red, AT8 monoclonal antibody) in the Hp, EC, TC and PFC of the three SMI Monkey 8, 9 and 10, and normal aged Monkey 1, 2 and 4, showing that in these AD related regions, the amount of AT8 positive cells in the SMI monkeys were obviously larger than that of normal aged monkeys. Scale bars: 200 μm in general view; 50 μm in larger view. (b-d) Quantitative comparisons of the percentage of the AT8 immunopositive areas of the SMI Monkey 8, 9 and 10 and that of the 3 normal aged monkeys in the Hp (Monkey 8, *P*=0.022; Monkey 9, *P*=0.071; Monkey 10, *P*=0.009), EC (Monkey 8, *P*=0.07; Monkey 9, *P*=0.046; Monkey 10, *P*=0.07), TC (Monkey 8, *P*=0.01; Monkey 9, *P*=0.438; Monkey 10, *P*=0.011) and PFC (Monkey 8, *P*=0.011; Monkey 9, *P*=0.026; Monkey 10, *P*=0.016), by one-sample t-test, respectively (for details, please see Methods). The results show that in most of the AD related regions the amount of AT8 immunopositive areas in the SMI monkeys is significantly larger than that of normal aged monkeys (n=3). The data of normal aged monkey group are represented as the mean±SEM. *: *P*<0.05, **: *P*<0.01.

**Figure S3.**
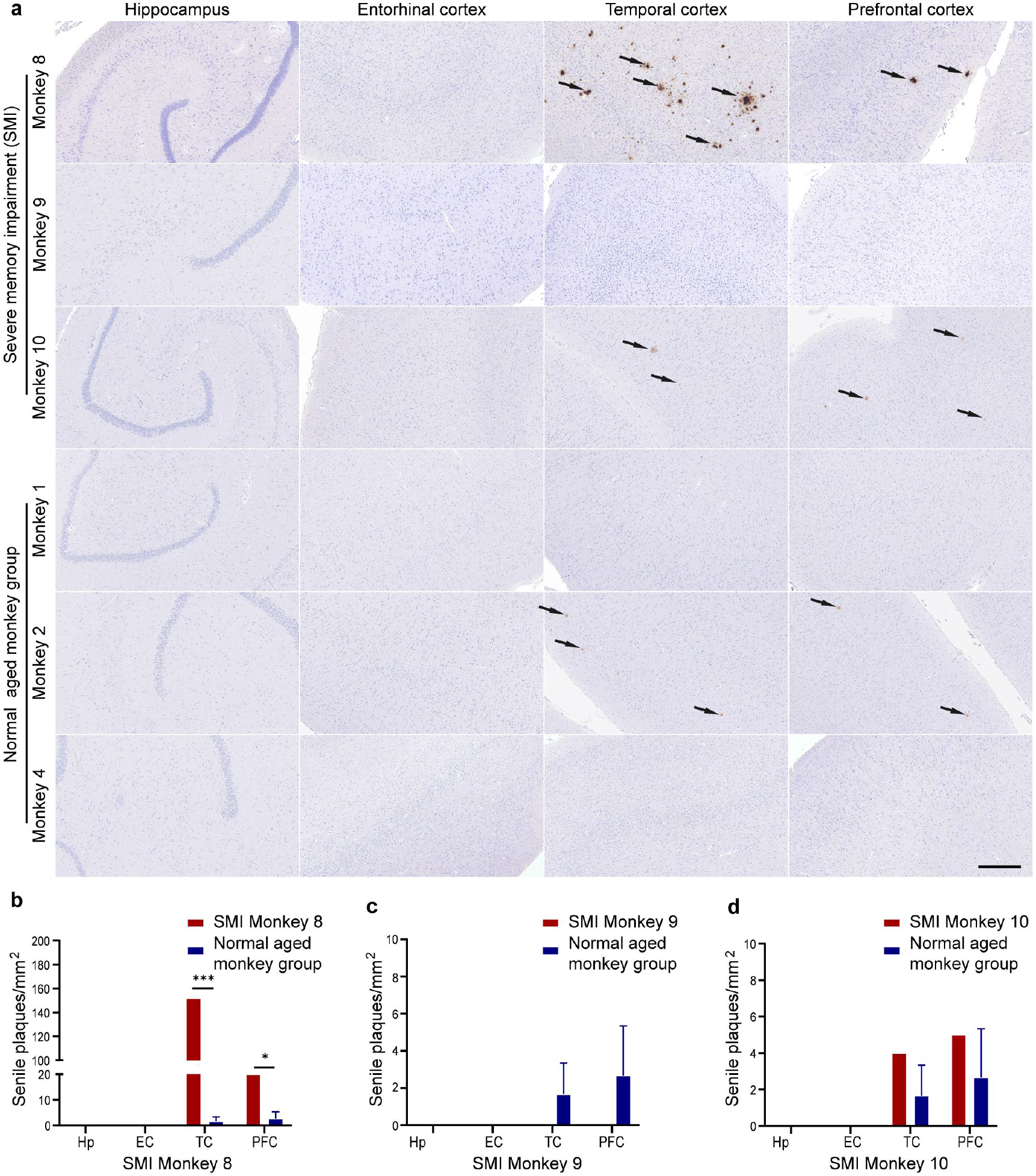
Pathological images and quantitative analyses of SPs in the brains of the three SMI monkeys and the three normal aged monkeys. (a) Representative images of SPs (brown, 6E10 monoclonal antibody, immunohistochemistry staining) in the EC, TC and PFC of the SMI Monkey 8, 9 and 10 and normal aged Monkey 1, 2 and 4, showing the presence of classical SPs in the AD related cortical areas of the SMI Monkey 8 and 10 and a small amount, less typical SPs in the normal aged Monkey 2. This is consistent with human data. Some autopsy studies report that some aged people have SPs or/and NFTs pathology in their brains with normal cognition. Black arrows mark the SPs in the TC and PFC of the SMI Monkey 8 and 10 and normal aged Monkey 2. Scale bar: 500 μm. (b-d) Comparison of the densities of the 6E10 immuno-positive SPs among the SMI Monkey 8, 9 and 10 and those of the normal aged monkeys (n=3) in the EC, TC (Monkey 8, *P*<0.001; Monkey 10, *P*=0.296) and PFC (Monkey 8, *P*=0.023; Monkey 10, *P*=0.474), by one-sample t-test, respectively (for details, please see Methods). The data of normal aged monkeys are represented as the mean ± SEM. *: *P*<0.05, ***: *P*<0.001.

**Figure S4.**
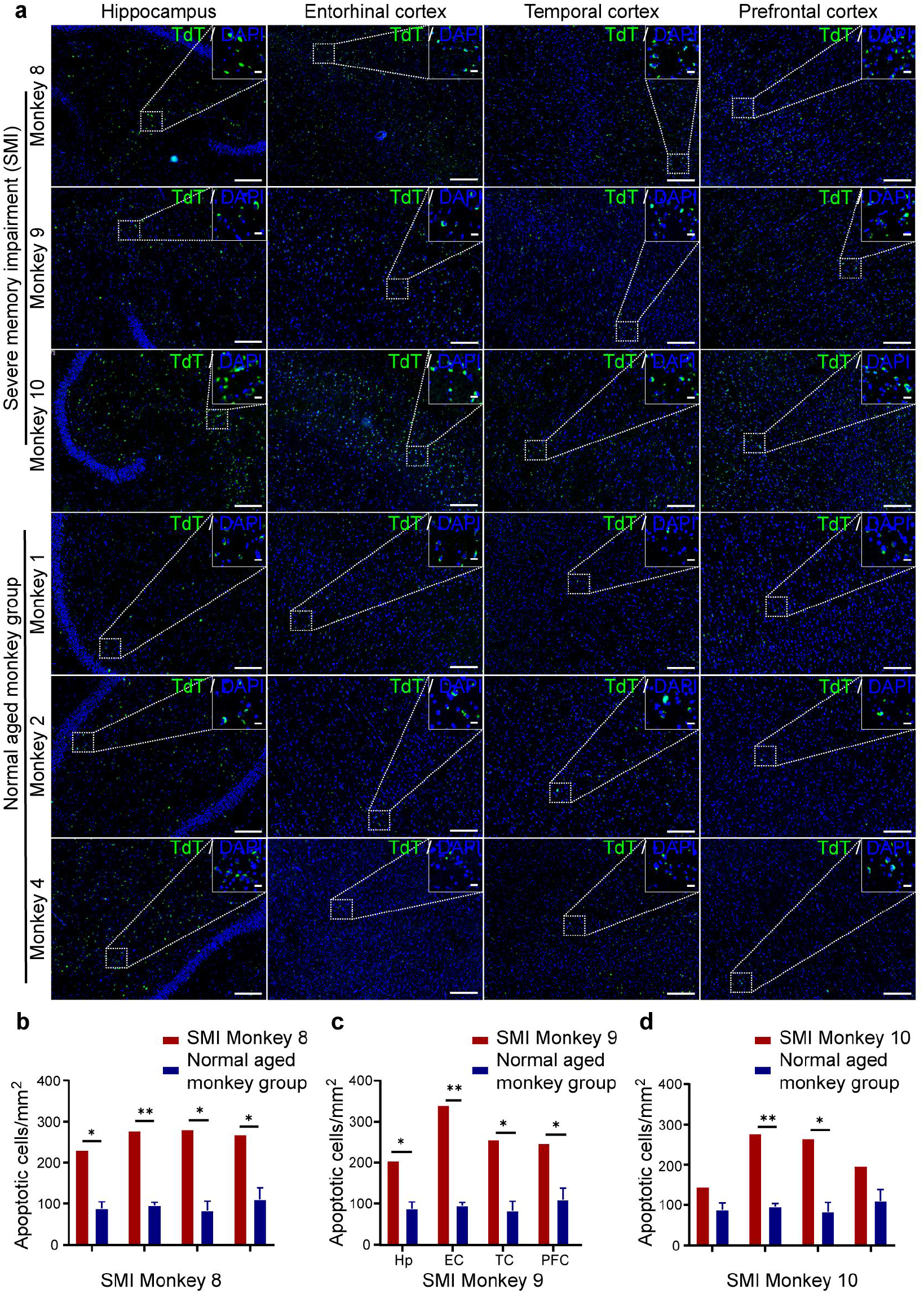
TdT labeling and quantitative analyses of apoptotic cells in the AD related cortical regions of the three SMI monkeys and three normal aged monkeys. (a) Representative images of TdT labeled apoptotic cells (green) in the Hp, EC, TC and PFC of the SMI Monkey 8, 9 and 10 and normal aged Monkey 1, 2 and 4, showing that in these AD related regions the number of TdT labeled apoptotic cells in the SMI monkey’s brain are obviously larger than those in the normal aged monkey’s brain. Scale bars: 200 μm in general view; 50 μm in larger view. (b-d) Quantitative comparison of the densities of apoptotic cells among the SMI Monkey 8, 9 and 10 and the 3 normal aged monkeys as a group in the Hp (Monkey 8, *P*=0.013; Monkey 9, *P*=0.019; Monkey 10, *P*=0.075), EC (Monkey 8, *P*=0.002; Monkey 9, *P*=0.001; Monkey 10, *P*=0.002), TC (Monkey 8, *P*=0.014; Monkey 9, *P*=0.017; Monkey 10, *P*=0.016) and PFC (Monkey 8, *P*=0.03; Monkey 9, *P*=0.038; Monkey 10, *P*=0.089), by one-sample t-test, respectively (for details, please see Methods). The results revealed that in most of these AD related regions the densities of TdT labeled apoptotic cells in the SMI monkeys were significantly higher than those in the normal aged monkey group (n=3). The data of normal aged controls are represented as the mean±SEM. *: *P*<0.05, **: *P*<0.01.

## Notes

### Competing Interest Statement

The authors have declared no competing interest.

