## Supplementary figures and images for "Naturally occurring Alzheimer’s disease in rhesus monkeys"

### Supplementary Figure S1

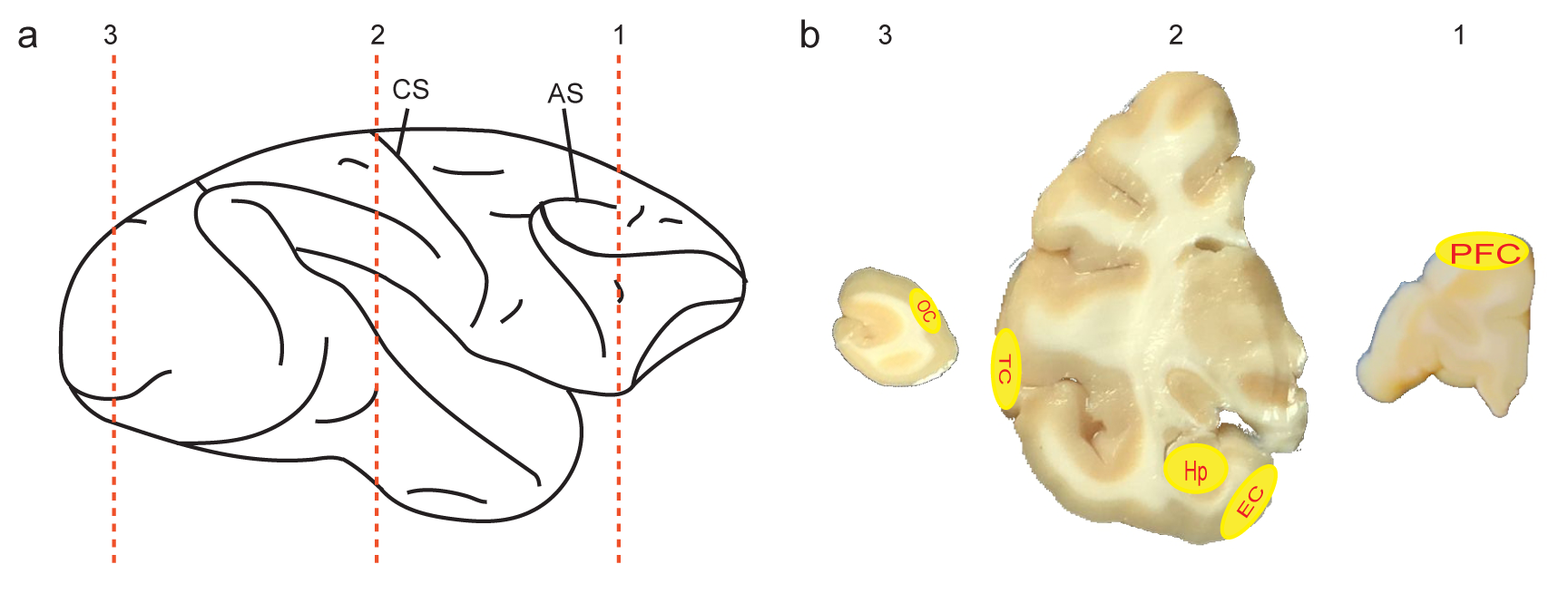

### Supplementary Figure S2

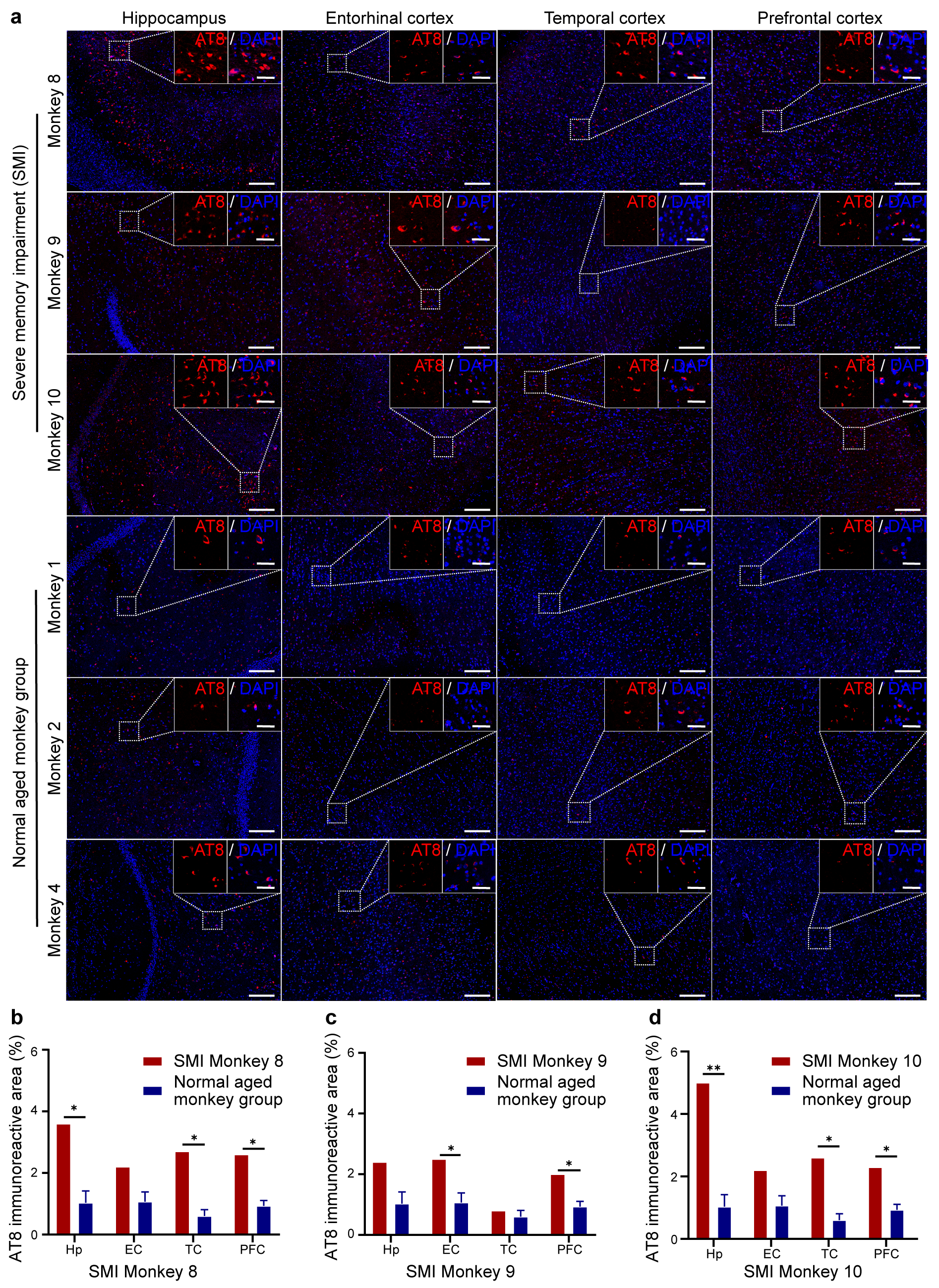

### Supplementary Figure S3

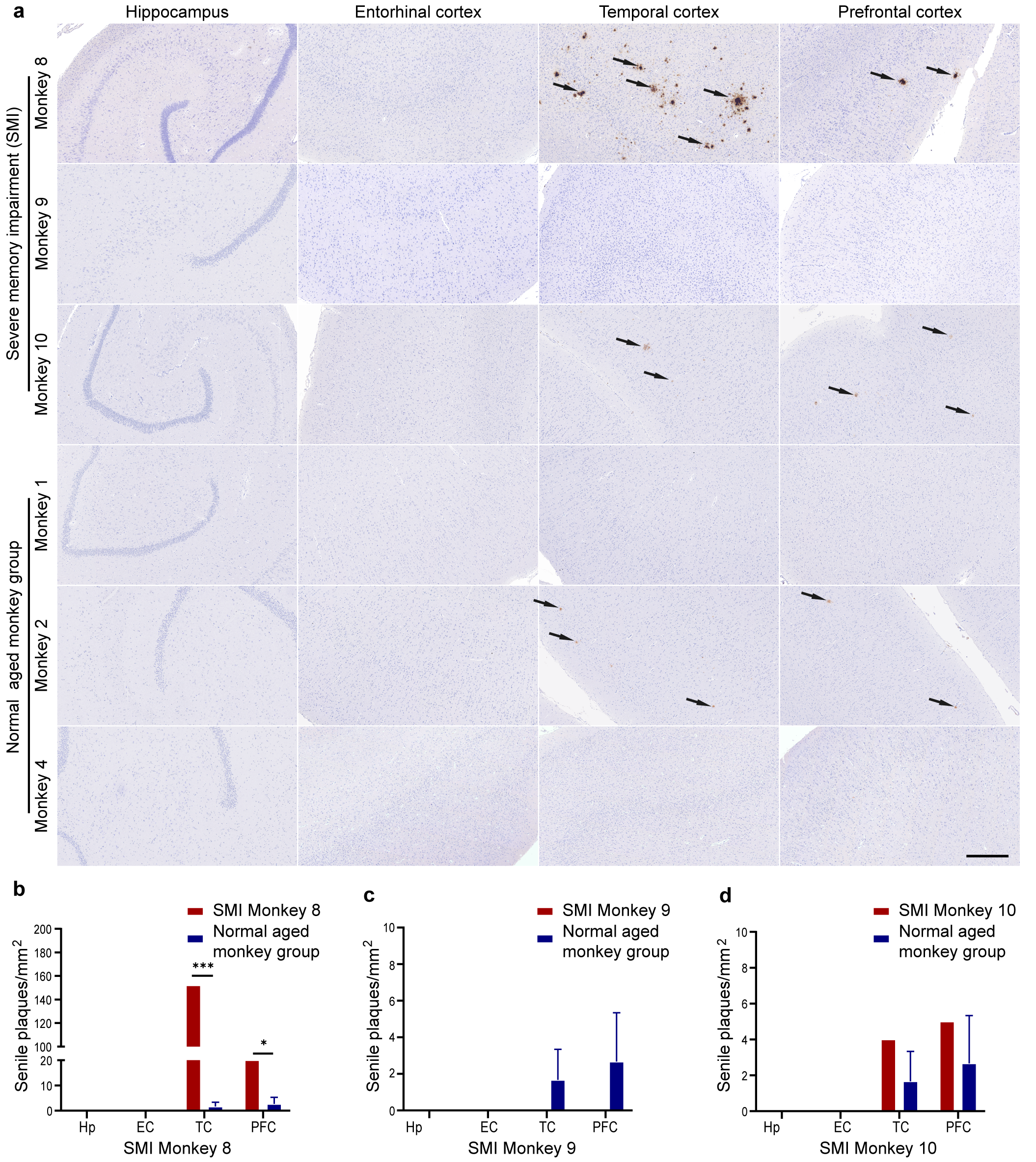

### Supplementary Figure S4

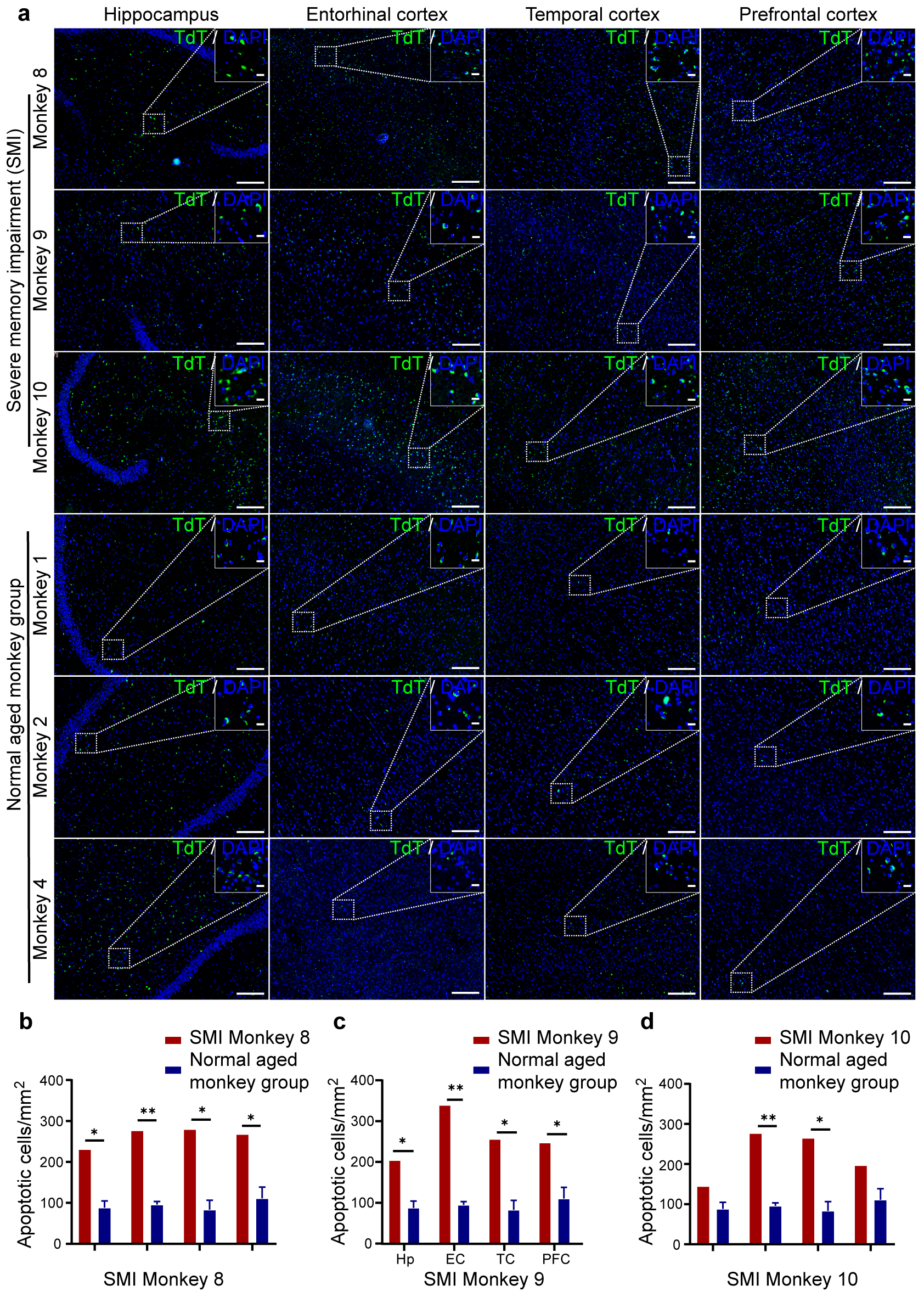
